# Level-Based Analysis of Genetic Algorithms and Other Search Processes

**DOI:** 10.1101/084335

**Authors:** Dogan Corus, Duc-Cuong Dang, Anton V. Eremeev, Per Kristian Lehre

## Abstract

Understanding how the time-complexity of evolutionary algorithms (EAs) depend on their parameter settings and characteristics of fitness landscapes is a fundamental problem in evolutionary computation. Most rigorous results were derived using a handful of key analytic techniques, including drift analysis. However, since few of these techniques apply effortlessly to population-based EAs, most time-complexity results concern simplified EAs, such as the (1 + 1) EA.

This paper describes the *level-based theorem*, a new technique tailored to population-based processes. It applies to any non-elitist process where o spring are sampled independently from a distribution depending only on the current population. Given conditions on this distribution, our technique provides upper bounds on the expected time until the process reaches a target state.

We demonstrate the technique on several pseudo-Boolean functions, the sorting problem, and approximation of optimal solutions in combinatorial optimisation. The conditions of the theorem are often straightforward to verify, even for Genetic Algorithms and Estimation of Distribution Algorithms which were considered highly non-trivial to analyse. Finally, we prove that the theorem is nearly optimal for the processes considered. Given the information the theorem requires about the process, a much tighter bound cannot be proved.

## 1 Introduction

The theoretical understanding of Evolutionary Algorithms (EAs) has advanced signi cantly over the last decade. A contributing factor for this success may have been the strategy to analyse simple settings before proceeding to more complex scenarios, while at the same time developing appropriate analytic techniques. In particular, much of the work assumed a population size of one, and no crossover operator. Current approaches to analysing evolutionary algorithms often rely on one or more of these simplifying assumptions.

This paper presents a general-purpose technique to analyse a large class of search heuristics involving non-overlapping populations. In our framework, each individual of the current population is independently sampled from the same distribution over the search space parametrised by the previous generation. A similar modelling of the search process first appeared in [56] to analyse Genetic Algorithms (GAs) however as far as we know, mainly results at the limit of infinite population were established. In this paper, we give the following general result for finite populations. Given some requirement on the upper tails of this distribution over an ordered partition of the search space and a minimum requirement on the population size, our method will guarantee an upper bound on the expected runtime to reach the last set of the partition.

Particularly, the partition of the search space is similar to the well-known fitness-level technique [58] to analyse *elitist* EAs, however at our general level of describing the search process, the traditional requirement on a fitness-based (this will be properly defined later on) partition is no longer required. Applications of the fitness-level technique itself are widely known in the literature for classical *elitist* EAs [58]. One of the first examples of using this technique in the analysis of non-elitist EAs is [24] where lower and upper bounds on the expected proportions of the population above certain fitness levels were found.

Related to our work, early research on analysing population-based EAs often ignored recombination operators. The family tree technique was introduced in [59] to analyse the (*μ* + 1) EA. The performance of the (*μ* + *μ*) EA for different settings of the population size was conducted in [33] using Markov chains to model the search processes, and in [5] using a similar argument to fitness-levels. The analysis of parallel EAs in [38] also made use of the fitness-levels argument. The ine ciency of standard fitness proportionate selection without scaling was shown in [46] and in [39] using drift analysis [30]. In the recently introduced switch analysis, the progress of the EA is analysed relative to an easier understand reference process [60]. When the method applies, bounds on the runtime of the reference process can be translated into bounds on the original process. In current applications of this method, the reference process is RLS^=^, a simple local search algorithm. It remains to be seen how such simple search heuristics can approximate the population dynamics of complex EAs.

Over the recent years, runtime analysis of EAs with recombination, often referred to as Genetic Algorithms, has been subject to increasing interest. Generalising the work in [46], [48,49] showed that the Simple Genetic Algorithm [56] is ine cient on OneMax, even when crossover is used. A long sequence of work has attempted to show that enabling crossover can reduce the runtime. It has been shown that adding crossover to the (*μ* + 1) EA can decrease the runtime on the Jump problem, however only for small crossover probabilities [34, 36]. For realistic crossover probabilities, it was shown that (*μ* + 1) GA can decrease the runtime by an exponential factor on instances of an FSM testing problem, however this result assumes a deterministic crowding diversity mechanism [41]. With the same setting on the standard OneMax function, crossover was shown to lead to a constant speedup in [54], however this result assumed a tailored selection mechanism. Seeking the construction of an e cient *unbiased* algorithms for OneMax, [20] introduced the (1+(λ,λ)) GA and showed a signi cant speed up with the right choices of the o spring population size [18,19]. Another modified GA, but this time *non-elitist*, was introduced in [51], and its e ciency was proved on the noisy version of OneMax function. In [43], a runtime result is proposed for a class of convex search algorithms, including some non-elitist GAs with gene pool recombination and no mutation, on the so-called *quasi-concave fitness landscapes*. As a corollary, it has been shown that the convex search algorithm has 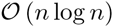 expected runtime on LeadingOnes. Those results gave the impression that adjustments or modifications to the standard setting of GAs, here elitist, are often required to illustrate the advantage of crossover. Until recently, it has been shown that the standard (*μ* + 1) GA without too low crossover probability has a speed up of Ω(*n*/log(*n*)) on the Jump problem com-pared to mutation-only algorithms [12].

Significant progress in developing and understanding a formal model of canonical GA and its generalisations was was made in [56] using dynamical systems. In particular it turned out that the behaviour of the dynamical systems model is closely related to the local optima structure of the problem in the case of binary search spaces [57]. Most of the findings in [56, 57] apply to the infinite population case, so it is not clear how these results can be used in runtime analysis of EA.

A relatively new paradigm in Evolutionary Computation is Estimation of Distribution Algorithm (EDA) [37]. Unlike traditional EAs which use explicit genetic operators such as mutation, recombination and selection, an EDA builds a probabilistic model for sampling new search points so that the probability of creating an optimal solution via sampling eventually increases high. The algorithm often starts with a specific probabilistic model, which is gradually updated through selected solutions of intermediate samplings. Over the recent years, many variants of EDAs have been proposed, along with theoretical investigations on their convergence and scalability, e. g. [29, 45, 50, 53, 61]. However, rigorous runtime analysis results for this particular class of algorithms on discrete domain are still sparse. The first analysis of this kind was conducted in [21] for the compact Genetic Algorithm (cGA) [31] on linear functions. Further work showed that this algorithm can be resilient to noise [27].

Another simple EDA is the Univariate Marginal Distribution Algorithm (UMDA) which was proposed in [44], and analysed in a series of papers [6–9]. With the *n*-dimensional Hamming cube as search space, each generation of UMDA consists of first sampling a population of solutions based on a vector (*p_i_*)_∈[*n*]_ of frequencies, i. e. assuming independence between bit positions, then summarising the selected solutions as the new sampling vector for the next generation. The initial result of [7] was provided for LeadingOnes and a harder function known as TrapLeadingOnes under the so-called “no-random-error” assumption and with a sufficiently large population. The assumption was lifted due to the technique presented in [9]. Nevertheless, the analysis assumes an unrealistically large population size, leading in overall to a too high bound on the expected runtime. Note also that there are two versions of the algorithm based on whether or not margins are imposed to *p_i_*, the difference between the two in terms of time complexity for various functions are discussed in [8]. More interestingly, [6] showed that UMDA without margins beats the (1 + 1) EA on a particular function called SubString. However, it is not recommended to use UMDA without margins in practice, as the algorithm can always end up with a premature convergence.

In this paper, we show that all non-elitist EAs with or without crossover, and even UMDA can be cast and analysed in the same framework. A preliminary version of the paper was communicated in [10]. This followed the line of work dated back to the introduction of a fitness-level technique to analyse *non-elitist* EAs with linear ranking selection [42], later on generalised to many selection mechanisms and *unary variation operators* [39], with a re ned result in [16]. The original fitness-level technique and its generalisation to the level-based technique have already found a number of applications, including analysis of EAs in uncertain environments, such as partial information [16], noisy fitness functions [14], and dynamic fitness functions [13]. It has also been applied to analyse the runtime of complex algorithms, such as GAs for shortest paths [11], EDAs [15], and self-adaptive EAs [17].

The present work improves the main result of [10] in many aspects. A more careful analysis of the population dynamics leads to a much tighter expression of the runtime bound compared to [10], immediately implying improved results in the previously mentioned applications. In particular, the leading term in the runtime is improved by a factor of Ω(*δ*^−3^), where characterises how fast good individuals can populate the population. This signi cantly improves the results of [14] and [16] concerning noisy optimisation, for which is often very small (e. g. 1/*n*). We also provide guideline how to use the theorem to analyse the runtime of non-elitist processes. Selected examples are given for the cases of GAs and UMDA in optimising standard pseudo-Boolean functions, a simple combinatorial problem, and in searching for local optima of NP-hard problems. Furthermore, we prove that the level-based theorem is close to optimal for the class of evolutionary processes it applies to.

The paper is structured as follows. In Section 2, we first present the general scheme of the algorithms covered by the main result of the paper and show how the GAs fit as special cases into this scheme. The main result of the paper is then presented along with a set of corollaries tailored to specific cases. Applications of the main result to difeerent GAs are considered in Section 4. The section starts with runtime analysis of the Simple Genetic Algorithm on standard functions, followed by the results for combinatorial optimisation problems, finally, the main theorem is again applied to analyse an Estimation of Distribution Algorithm. Section 6 considers the tightness of the level-based theorem. Finally, concluding remarks are given in Section 7.

## 2 Main result

### 2.1 Abstract algorithmic scheme

We consider population-based algorithms at a very abstract level in which fitness evaluations, selection and variation operations, which depending on the current population *P* of size λ, are represented by a distribution *D*(*P*) over a finite set *χ*. More precisely, the current population *P* is a vector (*P*(1),…, *P*(λ)) where *P*(*i*) ∈ *χ* for each *i* ∈ [λ]. *D* is a mapping from *χ*^λ^ into the space of probability distributions over *χ*. The next generation is obtained by sampling each new individual independently from *D*(*P*). This scheme is summarised in Algorithm 1. Here and below, for any positive integer *n*, we define [*n*]: = {1, 2,…, *n*}.

#### Algorithm 1

Population-based algorithm.

**Require:**

Finite state space *χ*, and population size 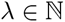,

Mapping D from *χ*^λ^ to the space of prob. dist. over *χ*.

Initial population *P*_0_ ∈ *χ*^λ^.

1: **for** *t* = 0, 1, 2,… until termination condition met **do**

2: Sample *P*_*t*+1_(*i*) ~ *D*(*P_t_*) independently for each *i* ∈ [λ]

3: **end for**

A scheme similar to Algorithm 1 was studied in [56], where it was called *Random Heuristic Search* with *an admissible transition rule*. Some examples of such algorithms are Simulated Annealing (more generally any algorithm with the population composed of a single individual), Stochastic Beam Search [56], Estimation of Distribution Algorithms such as the Univariate Marginal Distribution Algorithm [6] and the Genetic Algorithm [28]. The previous studies of the framework were often limited to some restricted settings [47] or mainly focused on infinite populations [56]. In this paper, we are interested in finite populations and develop a general method to deduce the expected runtime of the search processes defined in terms of *number of produced search points*. This can be translated to the *number of evaluations* once a specific algorithm is instantiated and the optimisation scenario is specified (e. g. see [16]).

We illustrate the general scheme of Algorithm 1 on the example of GA, which is Algorithm 2. The term Genetic Algorithm is often applied to EAs that use recombination operators with some a priori chosen probability *p*_c_ > 0. Here the standard operators of GA are formally represented by transition matrices:

#### Algorithm 2

Genetic algorithm.

**Require:**

Finite state space *χ*,

Operators: Sel, Cross and Mut,

Population size 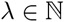 and recombination rate *p*_c_ ∈ [0, 1].

1: *P*_0_ ~ Unif(*χ*^λ^)

2: **for** *t* = 0, 1, 2,… until termination condition met **do**

3: **for** *i* = 1 to λ **do**

4: *u*: = *P_t_*(Sel(*P_t_*)), *v*: = *P_t_*(Sel(*P_t_*)).

5: 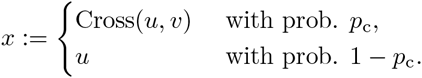

6: *P*_*t* + 1_(*i*): = Mut(*x*).

7: **end for**

8: **end for**

- *p*_sel_: [λ] *χ*^λ^ → [0, 1] represents selection operator Sel: *χ*^λ^ → [λ] which is randomised, where *p*_sel_(*i*|*P_t_*) is the probability of selecting the i-th individual from population *P_t_*. This probability can depend on the search point *P_t_*(*i*), its relationship to the other search points in *P_t_*, and their mappings to fitness values by a function 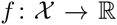 that the algorithm aims to optimise. Throughout the paper, we assume w.l.o.g. the maximisation of *f*.
- *p*_mut_: *χ* × *χ* → [0, 1], where *p*_mut_(*y*|*x*) is the probability of mutating *x* ∈ *χ* into *y* ∈ *χ* by a randomised mutation operator Mut: *χ* → *χ*.
- *p*_xor_: *χ*^2^ × *χ*^2^ → [0, 1], where *p*_xor_(*x*|*u*, *v*) is the probability of obtaining *x* as a result of randomised crossover operator (or recombination) between *u*, *v* ∈ *χ*. In what follows, crossover is denoted by Cross: *χ* × *χ* → χ.

Clearly, conditioned on the current population *P_t_*, each individual *P*_*t* + 1_(*i*) of the next generation is independently sampled from the same distribution which is parametrised by *P_t_*. Thus, lines 4-6 of Algorithm 2 can be summarised as *P*_*t* + 1_(*i*) ∼ *D*(*P_t_*) for some *D* induced by the genetic operators Sel, Cross and Mut. The algorithm fits perfectly in the scheme of Algorithm 1.

### 2.2 Level-based theorem

This section states the main result of the paper, a general technique for obtaining upper bounds on the expected runtime of any process that can be described in the form of Algorithm 1. We use the following notation. The natural logarithm is denoted by ln(·). Suppose that for some *m* there is an ordered partition of *χ* into subsets (*A*_1_,…, *A_m_*) called levels, we define 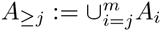, i. e. the union of all levels above level *j*. An example of a partition is the *canonical* partition, where each level regroups solutions having the same fitness value (see e.g. [39]). This partition is classified as *fitness-based* or *f*-based, if *f*(*x*) < *f*(*y*) for all *x* ∈ *A_j_*, *y* ∈ *A*_*j* + 1_ and all *j* ∈ [*m* − 1]. As a result of the algorithmic abstraction, our main theorem is not limited to this particular type of partition. Let *P* ∈ *χ*^λ^ be a population vector of a finite number 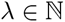 of individuals. Given any subset *A* ⊆ *χ*, we write |*P* ∩ *A*|: = |{*i* | *P*(*i*) ∈ *A*}| to denote the number of individuals in population *P* that belong to the subset *A*.

#### Theorem 1.

Given a partition (*A*_1_,…, *A_m_*) of *χ*, define *T*: = min{*tλ* | |*P_t_* ∩ *A_m_*| > 0} to be the first point in time that elements of *A_m_* appear in *P_t_* of Algorithm 1. If there exist *z_1_*,…, *z_m−1_*; *δ* ∈ (0, 1], and *γ*_0_ ∈ (0, 1) such that for any population *P* ∈ *χ^λ^*,

**(G1)** for each level *j* ∈ [*m* − 1], if |*P* ∩ *A_≥j_*| ≥ γ_0_λ then

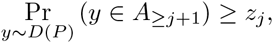

**(G2)** for each level *j* ∈ [*m* − 2], and all *γ* ∈ (0, γ_0_] if |*P* ∩ *A_≥j_*|≥ γ_0_λ and |*P* ∩ *A_≥j°1_*| ≥ γ_0_λ then

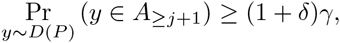

**(G3)** and the population size 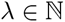 satisfies

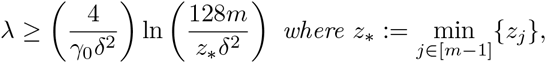
then

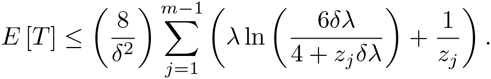

Informally, the two first conditions require a relationship between the current population P and the distribution *D*(*P*) of the individuals in the next generation: Condition (G1) demands that the probability of creating an individual at level *j* + 1 or higher is at least *z_j_* when some fixed portion γ_0_ of the population has reached level *j* or higher. Furthermore, if the number of individuals at level *j* + 1 or higher is at least γλ > 0, condition (G2) requires that their number tends to increase further, e.g. by a multiplicative factor of 1 + δ. Finally, (G3) requires a sufficiently large population size. When all conditions are satisfied, an upper bound on the expected time for the algorithm to create an individual in *A_m_* can be guaranteed.

We suggest to follow the five steps below when applying the level-based theorem.

1. Identify a partitioning of the search space which re ects the “typical” progress of the population towards the target set *A_m_*.
2. Find parameter settings of the algorithm and corresponding parameters γ_0_ and *δ* of the theorem, such that condition (G2) can be satisfied. It may be necessary to adjust the partitioning of the search space.
3. For each level *j* ∈ [*m* − 1], estimate lower bounds *z_j_* such that condition (G1) holds.
4. Determine the lower bound on the population size λ in (G3) using the parameters obtained in the previous steps.
5. Once all conditions are satisfied, compute the bound on the expected time from the conclusion of the theorem. A simple way (not necessarily the only way) to evaluate the sum 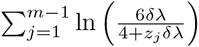 is to underestimate the denominator 4 + *z_j_δ*λ in each term by either 4, or *z_j_δ*λ. This gives the bounds *m* ln(3λ/2), or 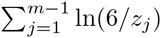 for this sum.

Some iterations of the above steps may be required to find parameter settings that yield the best possible bound.

We now illustrate this methodology on a simple example.

#### Corollary 2

For any *λ* ≥ 72(ln(n) + 9), the expected number of points created until the population of Algorithm 3 contains the point *n* is 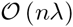.

#### Algorithm 3

Example algorithm to illustrate Theorem 1.

**Require:** Finite state space *χ* = {1,…, *n*} for some *n* ∈ N.

1: *P*_0_ ~ Unif(*χ*^λ^), i.e. initial population sampled u.a.r.

2: **for** *t* = 0, 1, 2,… until termination condition met **do**

3: **for** *i* = 1 to λ **do**

4: Sort the current population *P_t_* = (*x*_1_,…, *x*_λ_) such that *x*_1_ ≥ *x*_2_ ≥ *x*λ.

5: *z*: = *x_k_* where *k* ~ Unif({1,…, *χ*/2}).

6: *y*: = *z* + Unif({−*c*, 0, 1}) for any *c* ∈ [*n*]

7: *P*_*t* + 1_(*i*): = max{1, min{*y*, *n*}}.

8: **end for**

9: **end for**

The purpose of Algorithm 3 is to illustrate the application of Theorem 1 on a very simple example. The search space *χ* is the set of natural numbers between 1 and *n*. A population is a vector of λ such numbers, and the implicit objective is to obtain a population containing the number *n*. Following the scheme of Algorithm 1, the operator *D* corresponds to lines 4-6. The new individual *y* is obtained by first selecting uniformly at random one of the best λ/2 individuals in the population (lines 4 and 5), and “mutating” this individual by adding 1, subtracting *c*, or do nothing, with equal probabilities. The value of *c* does not matter in our analysis. Note that for *c* being fixed to 1, one could ignore the selection steps and easily come up with a rough bound 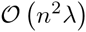. However, for other choices of *c*, e. g. equal to *n* or randomly picked, without the tool proposed in Theorem 1 it is much less obvious how such a process should be approached and analysed.

We now carry out the steps described previously.

Step 1: It seems natural to partition the search space into *m* = *n* levels, where *A_j_*: = {*j*} for all *j* ∈ [*m*].

Step 2: Assume that the *current level* is *j* < *n* − 1. This means that in *P_t_*, there are γ_0_λ individuals in *A*_≥*j*_, i.e. with fitness at least *j*, and at least γλ but less than γ_0_λ individuals in *A*_≥*j* + 1_, i.e. with with fitness at least *j* + 1. We need to estimate Pr_*y*~*D*(P_t_)_(*y* ∈ *A*_≥*j*+1_), i.e., the probability of producing an individual with fitness at least *j* + 1. To this end, we say that a selection event is “good” if in step 5, the algorithm selects an individual in *A*_≥*j*+1_, i.e. with fitness at least *j* + 1. If γ ≤ 1/2, then the probability of a good selection event is at least γλ/(λ/2) = 2γ. And we say that a mutation event is “good” if in line 5, the algorithm does not subtract 1 from the selected search point. The probability of a good mutation event is 2/3. Selection and mutation are independent events, hence we have shown for all γ ∈ (0, 1/2] that

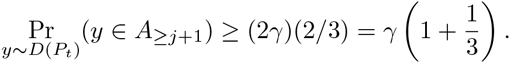

Condition (G2) is therefore satisfied with *δ* = 1/3 if we choose any positive constant γ_0_ ≤ 1/2.

Step 3: Assume that population *P_t_* has at least γ_0_λ individuals in *A*_≥*j*_. In this case, the algorithm produces an individual in *A*_≥*j*+1_ if in line 5 it selects an individual in *A*_≥*j*_ and mutates the individual by adding 1 in line 6. If we now fix γ_0_ = 1/2, the probability of selecting an individual in *A*_≥*j*_ is 1 by the assumption. Furthermore, the probability of adding 1 to the selected individual is exactly 1/3. Hence, we have shown

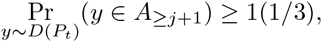

and we can satisfy condition (G1) by defining *z_j_*: = 1/3 for all *j* ∈ [*m* − 1].

Step 4: For the parameters we have chosen, it is easy to see by numerical calculation that the population size λ ≥ 72(ln(*n*) + 9) satis es condition (G3).

Step 5: We use 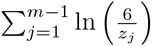 instead of 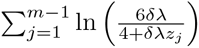, thus the expected time until the population has found the point *n* is no more than

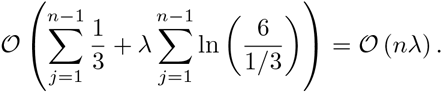

### 2.3 Proof of the level-based theorem

Theorem 1 will be proved using drift analysis, which is a standard tool in theory of randomised search heuristics. Our distance function takes into account both the “current level” of the population, as well as the distribution of the population around the current level. In particular, let the current level *Y_t_* be the highest level *j* ∈ [*m*] such that there are at least γ_0_λ individuals at level *j* or higher. Furthermore, for any level *j* ∈ [*m*], let 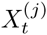 be the number of individuals at level *j* or higher. Hence, we describe the dynamics of the population by *m* + 1 stochastic processes 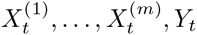, *Y_t_*. Assuming that these processes are adapted to a filtration 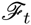, we write 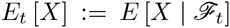 and 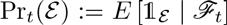. Our approach is to measure the distance of the population at time *t* by a scalar 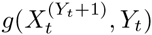, where *g* is a function that satis es the conditions in Definition 3.

#### Definition 3.

A function 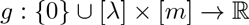 is called a level function if

1. ∀*x* ∈ {0} ∪ [λ], ∀*y* ∈ [*m* − 1] *g*(*x*, *y*) ≥ *g*(*x*, *y* + 1),
2. ∀*x* ∈ {0} ∪ [λ − 1], ∀*y* ∈ [*m*] *g*(*x*, *y*) ≥ *g*(*x* + 1, *y*), *and*
3. ∀*y* ∈ [*m* − 1] *g*(λ, *y*) ≥ *g*(0, *y* + 1).

It is clear from the definition that the sum of two level functions is also a level function. In addition, the three conditions ensure that the distance 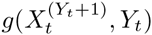 of the population decreases monotonically with the current level *Y_t_*. As the following lemma shows, this monotonicity allows an upper bound on the distance in the next generation which is partly independent of the change in current level.

#### Lemma 4.

If *Y_t+1_Y_t_*, then for any level function *g*

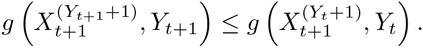

*Proof*. The statement is trivially true when *Y_t_* = *Y*_*t*+1_. On the other hand, if *Y*_*t*+1_ > *Y_t_*, then the conditions in Definition 3 imply

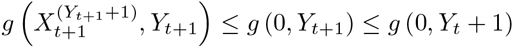

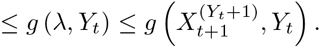

We can now give the formal proof of Theorem 1.

*Proof of Theorem 1*. We will prove the theorem using Lemma 22 (the additive drift theorem) with respect to the parameter *a* = 0 and a stochastic process

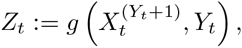

where *g* is a level-function to be defined, and 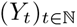 and 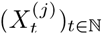 for *j* ∈ [*m*] are stochastic processes to be defined. We consider the filtration 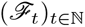 induced by the sequence of populations 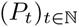.

We will assume w.l.o.g. that condition (G2) is also satisfied for *j* = *m* − 1, for the following reason. Given an Algorithm 1 with certain mapping *D*, consider Algorithm 1 with a difeerent mapping *D*′(*P*): If |*P* ∩ *A_m_*| = 0 then *D*′(*P*) = *D*(*P*); otherwise *D*′(*P*) assigns probability mass 1 to some element *x* of *P* that is in *A_m_*, e.g. to the first one among such elements. Note that *D*′ meets conditions (G1) and (G2). Moreover, (G2) holds for *j* = *m* − 1. For the sequence of populations 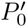, 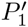,… of Algorithm 1 with mapping *D*′ we can put 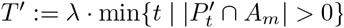. Executions of the original algorithm and the modified one before generation *T*′/λ are identical. On generation *T*′/λ both algorithms place elements of *A_m_* into the population for the first time. Thus, *T*′ and *T* are equal in every realisation and their expected values is the same.

For any level *j* ∈ [*m*] and time *t* ≥ 0, let the random variable

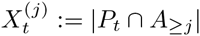

denote the number of individuals in levels *A*_≥*j*_ at time *t*. Because *A*_≥*j*_ is partitioned into disjoint sets *A_j_* and *A*_*j*+1_, the definition implies

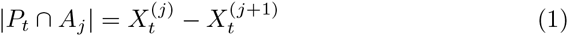

Algorithm 1 samples all individuals in generation *t* + 1 independently from distribution *D*(*P_t_*). Therefore, given the current population 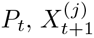 is binomially distributed

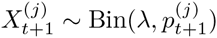

where 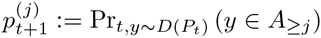 is the probability of sampling an individual in level *j* or higher.

The *current level Y_t_* of the population at time *t* ≥ 0 is defined as

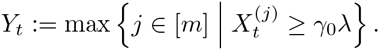

Note that 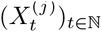 and 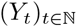 are adapted to the filtration 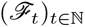 because they are defined in terms of the population process 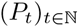.

When *Y_t_* < *m*, there exists a unique γ < γ_0_ such that

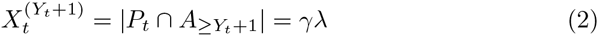

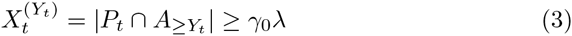

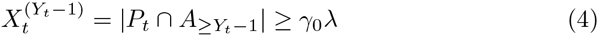

In the case of 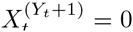, it follows from (1), (2) and (3) that |*P* ∩ *A_j_* = 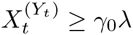. Condition (G1) for level *j* = *Y_t_* then gives

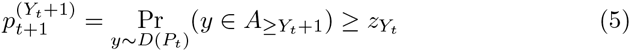

Otherwise if 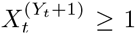, conditions (G1) and (G2) for level *j* = *Y_t_* with (2) and (3) imply

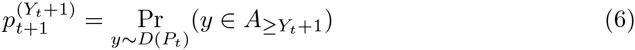

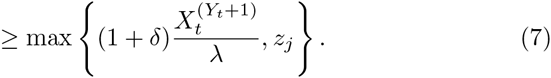

Condition (G2) for level *j* = *Y*_*t* − 1_ along with (3) and (4) also gives

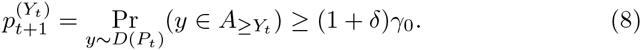

We now define the process 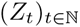 as *Z_t_*: = 0 if *Y_t_* = *m*, and otherwise, if *Y_t_* < *m*, we let 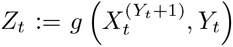, where for all *k*, and for all 1 ≤ *j* < *m*, *g*(*k*, *j*) = *g*_1_(*k*, *j*) + *g*_2_(*k*, *j*) and

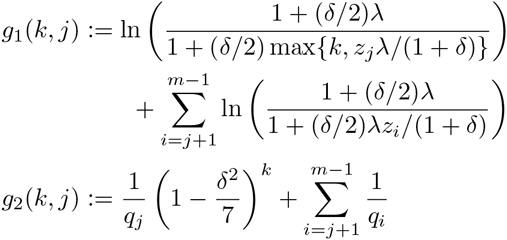

where *q_j_*: = 1 − (1 − *z_j_*)^λ^.

It follows from Lemma 21 that *g*(*k*, *j*) is a level function. Furthermore, *g*(*k, j*) ≥ 0 for all *k* ∈ {0} ∪ [λ] and all *j* ∈ [*m*]. Due to properties 1 and 2 of level functions and Lemma 18, the distance is always bounded from above by

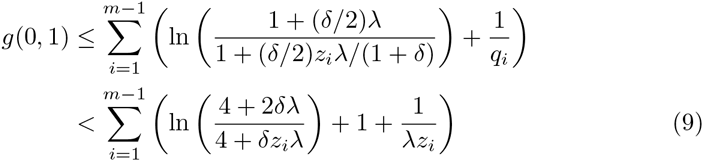

using that *z_i_* ≤ 1,

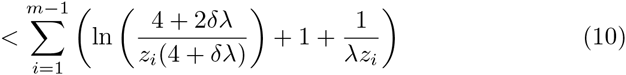

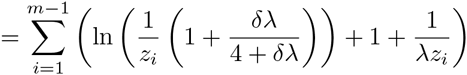

ln (*x*) ≤ *x* − 1 for all *x* > 0 so

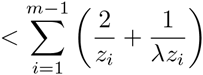

and *z_i_ z*_*_ and λ ≥ ⌈4 ln(128)⌉ = 20 from (G3) so

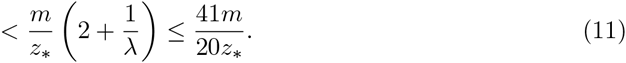

Hence, we have 0 ≤ *Z_t_* < *g*(0, 1) < ∞ for all 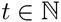 which implies that *E* [*Z_t_*] < ∞ for all 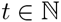, and condition 2 of the drift theorem is satisfied.

The “drift” of the process is the random variable

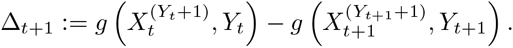

To compute the expected drift, we apply the law of total probability

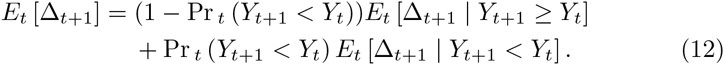

The event *Y*_*t* + 1_ < *Y_t_* holds if and only if 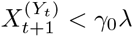. Due to (8), we obtain the following by a Cherno bound

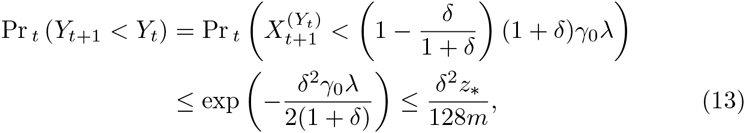

where the second last inequality takes into account the population size required by condition (G3). Given the low probability of the event *Y*_*t* + 1_ < *Y_t_*, it suffices to use the pessimistic bound from (11)

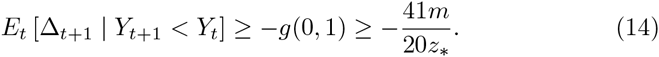

If *Y*_*t* + 1_ *Y_t_*, we can apply Lemma 4

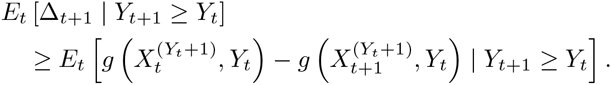

Note that event *Y*_*t* + 1_ ≥ *Y_t_* is equivalent to having 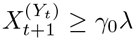, then due to Lemma 27, in the following we can skip the condition on the event when needed.

If 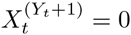, then 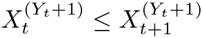 and

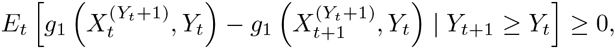

because the function *g*_1_ satisfies property 2 in Definition 3. Furthermore, we have the lower bound

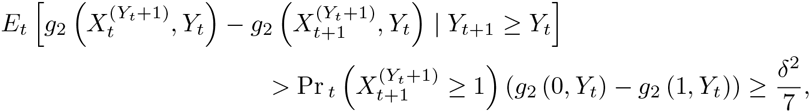

where the last inequality follows because of (5) and 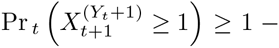 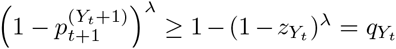, and *g*_2_ (0, *Y_t_*) − *g*_2_(1, *Y_t_*) = (1/*qY_t_*) = (*δ*^2^/7).

In the other case, where 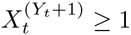, we obtain

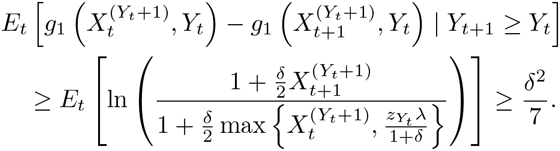

where the last inequality follows from (7) and Lemma 23. For function *g*_2_, we get

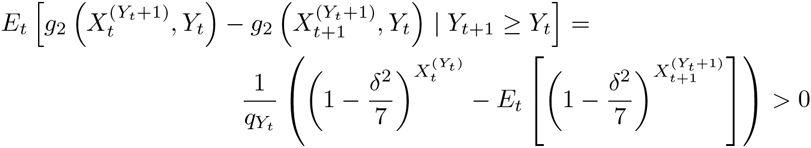

where the last inequality is due to Lemma 26, applied to

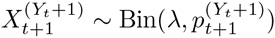

with 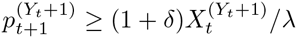 (see (7)) and the parameter

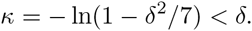

Taking into account all cases, we have

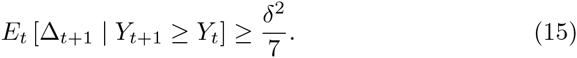

We now have bounds for all the quantities in (12) with (13), (14) and (15), and we get

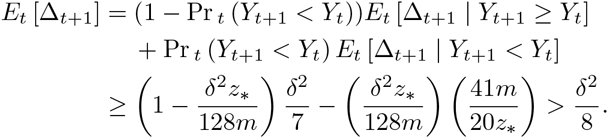

We now verify condition 3 of the drift theorem (Lemma 22), i.e., that *T* has finite expectation. Let *p*_*_: = min{(1 + *δ*)/ *δ z*_*_}, and note by conditions (G1) and (G2) that the current level increases by at least one with probability Pr_*t*_(*Y*_*t* + 1_ ≥ *Y_t_*) ≥ (*p*_*_)^γ_0_λ^. Furthermore, due to the definition of the modified process *D*′, if *Y_t_* = *m* then *Y*_*t* + 1_ = *m*. Hence, the probability of reaching *Y_t_* = *m* is lower bounded by the probability of the event that the current level increases in all of at most *m* consecutive generations, i.e., Pr*_t_*(*Y*_*t* + *m*_ = *m*) ≥ (*p*_*_)^γ_0_λ*m*^ > 0. It follows that *E*[*T*] < ∞.

By Lemma 22 and the upper bound on *g*(0, 1) in (9),

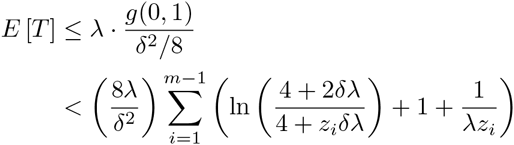

then using that 4 ≤ *δ*λ/5 from (G3) and (1/5 + 2)*e* < 6

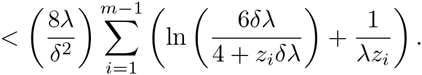

## 3 Tools for Analysis of Genetic Algorithms

This section provides two corollaries of Theorem 1 tailored to Algorithm 2. After that, we give sufficient conditions for tunable parameters of many selection mechanisms allowing the applicability of the corollaries.

Since no explicit fitness function is defined in Algorithm 2, no assumption on a *f*-based partition will be required by the corollaries. Nevertheless, we have to generalise the *cumulative selection probability* function of Sel operator, denoted *β*(γ, *P*) [16] which is defined relative to the fitness function *f*, to the one that is relative to the order of the partition (*A*_1_,…, *A_m_*).

Recall that to define *β*(γ, *P*) of Sel w. r. t. *f* for a population *P* of λ search points, we first assume (*f*_1_,…, *f*) to be the vector of sorted fitness values of *P*, i. e. *f_i_* ≥ *f*_*i* + 1_ for each *i* ∈ [λ − 1]. Then given γ ∈ (0, 1], define

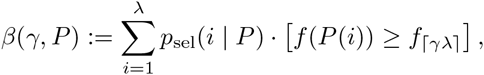

where [·] denotes the Iverson bracket.

Similarly, given a partition (*A*_1_,…, *A_m_*), if we use (*ℓ*_1_,…, *ℓ*_λ_) to denote the sorted levels of search points in *P*, i.e. *ℓ_i_* ≥ *ℓ*_*i* + 1_ for each *i* ∈ [λ − 1], then the cumulative *selection probability* of function of Sel w. r. t. (*A*_1_,…, *A_m_*) is

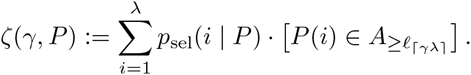

Let (*A*_1_,…, *A_m_*) be a f-based partition of *χ*, it follows from the above definitions that

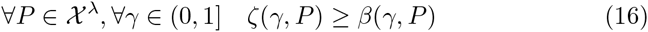

for which the equality occurs when the partition is canonical.

### 3.1 Analysis of non-permanent use of crossover

We first derive from Theorem 1 a corollary that is adapted to Algorithm 2 with *p_c_* < 1. This setting covers the case *p_c_* = 0, i. e. only unary variation operators are used. This specific case is the main subject of [16], and to some extent our corollary shares many similarities with the main theorem of that paper. As we will see later on, stronger and more general results can be claimed with the corollary.

#### Corollary 5.

Let (*A*_1_,…, *A*_m_) be a partition of *χ*, define *T*: = min {*tλ* | |*P* ∩ *A_m_*| > 0} to be the first point in time that Algorithm 2 with *p_c_* < 1 obtains an element of *A_m_*. If there exist *s*_1_,…, *s*_*m*− 1_, *s*_*_, *p*_0_, δ ∈ (0, 1], and a constant γ_0_ ∈ (0, 1) such that

**(M1)** for each level *j* ∈ [*m* − 1]

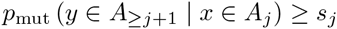

**(M2)** for each level *j* ∈ [*m* − 1]

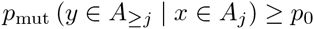

**(M3)** for any population *P* ∈ (χ \ *A_m_*) and ∈ (0, γ_0_]

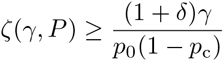

**(M4)** the population size λ satisfies

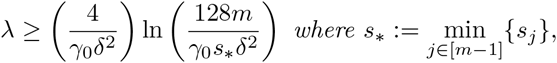
then

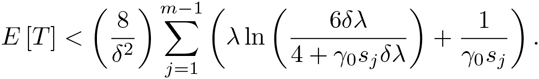

*Proof*. Following the guideline, we apply Theorem 1. Step 1 is skipped because we already have the partition. Step 2: We assume |*P* ∩ *A*_≥*j*_| ≥ γ_0_λ and |*P* ∩ *A*_≥*j*+1_| ≥ γλ > 0 for some γ ≤ γ_0_. Hence, to create an individual in *A*_≥*j*+1_ it suffices to pick an *x* ∈ |*P* ∩ *A_k_*| for any *k* ≥ *j* + 1 and mutate it to an individual in *A*_≥*k*_, the probability of such an event according to (M2) and (M3) is at least (1 − *p_c_*)ζ(γ, *P*)*p*_0_ ≥ (1 + *δ*)γ. So (G2) holds for the same *p*_0_, *δ* and γ_0_ as in (M3).

Step 3: We are given |*P* ∩ *A_j_*| ≥ ³_0_λ. Thus, with probability ζ(γ_0_, *P*), the selection mechanism chooses an individual *x* in either *A_j_* or *A*_≥*j*+1_. If the individual *x* belongs to *A_j_*, then the mutation operator will by condition (M1) upgrade the individual to *A*_≥*j*+1_ with probability *s_j_*. If the individual belongs to *A*_≥*j*+1_, then by (M2), the mutation operator maintains the individual in *A*_≥*j*+1_ with probability *p*_0_. Finally, no crossover occurs with probability 1 − *p_c_*. Hence, the probability of producing an individual in *A*_≥*j*+1_ is at least

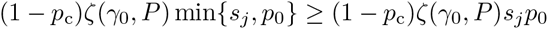

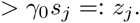

Thus (G1) holds for that choice of *z_j_* and *z*_*_: = γ_0_*s*_*_.

Step 4: Given our choice of *z*_*_, we have that condition (M4) implies condition (G3).

For the last step, all conditions (G1-3) are satisfied, and Theorem 1 gives

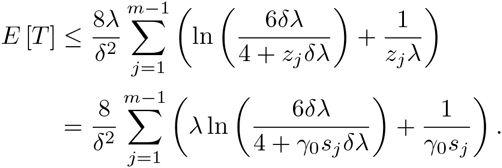

From the proof, we remark that any operator can be used in place of crossover in line 5 of Algorithm 2, and the result still holds. Therefore, the corollary is in fact applicable to a wider range of algorithms than just Algorithm 2.

### 3.2 Analysis of permanent use of crossover

The following corollary adapts Theorem 1 to the setting of Algorithm 2 with *p_c_* = 1.

#### Corollary 6.

Given a partition (*A*_1_,…, *A_m_*) of χ, let *T*: = min{*t*λ | |*P* ∩ *A_m_*| > 0} be the first point in time that Algorithm 2 with *p_c_* = 1 obtains an element of *A_m_*. If there exist *s*_1_,…, *s*_*m* − 1_, *s*_*_, *p*_0_, ε, ∈ (0, 1], and a constant γ_0_ ∈ (0, 1) such that

**(C1)** for each level *j* ∈ [*m* − 1]

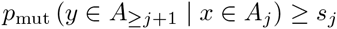

**(C2)** for each level *j* ∈ [*m*]

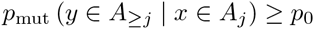

**(C3)** for each level *j* ∈ [*m* − 2]

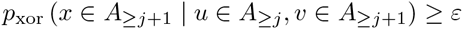

**(C4)** for any population *P* ∈ (χ \ *A_m_*)λ and γ ∈ (0, γ_0_]

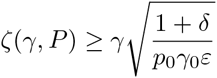

**(C5)** the population size satisfies

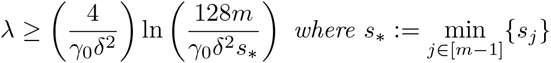
then

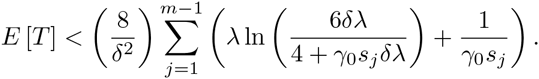

*Proof*. We apply Theorem 1 following the guideline. Again, Step 1 is skipped because the partition is already defined.

Step 2: We are given |*P* ∩ *A*_≥*j*_| ≥ γ_0_λ and |*P* ∩ *A*_≥*j*+1_| ≥ γλ > 0. To create an individual in *A*_≥*j*+1_, it suffices to pick the individual *u* in *A*_≥*j*_ and *v* in *A*_*j*+1_, then to produce an individual in *A_k_* for any *k* ≥ *j* + 1 by crossover and not destroy the produced individual by mutation. The probability of such an event according to (C2), (C3) and (C4) is bounded from below by ζ(γ_0_, *P*)ζ(γ, *P*)_ε*p*_0__ ≥ (1 + *δ*)γ. Condition (G2) is then satisfied with the same γ_0_ and *δ* as in (C4).

Step 3: We assume |*P* ∩ *A_j_*| ≥ γ_0_λ. Note that condition (C3) written for level *j* − 1 is *p*_xor_(*x* ∈ *A*_≥*j*_ | *u* ∈ *A*_≥*j* − 1_, *v* ∈ *A*_≥*j*_) ≥ ε, and because *A*_≥*j*_ ⊂ *A*_≥*j* − 1_ then *p*_xor_(*x* ∈ *A*_≥*j*_ | *u* ∈ *A*_≥*j*_, *v* ∈ *A*_≥*j*_) ≥ ε. To create an individual in *A*_≥*j*+1_, it then suffices to pick both u and *v* from *A*_≥*j*_ in line 4, then to produce an individual in *A_k_* for any *k* ≥ *j* by crossover, now if *k* = *j* we need to improve the produced individual by mutation, i. e. relying on (C1), otherwise if *k* > *j* it suffices not to destroy the produced individual by mutation, i. e. relying on (C2). It then follows from (C4) that the probability of producing an individual in *A*_≥*j*+1_ is at least

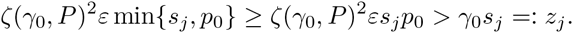

Condition (G1) then holds for that choice of *z_j_* and *z*_*_: = γ_0_*s*_*_.

Step 4: It follows from the above definition of *z*_*_ that (C5) implies (G3). In the last step, since all conditions (G1-3) are satisfied, Theorem 1 guarantees that

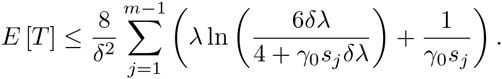

The corollary shares many similarities with Corollary 5, except that condition (C2) has to additionally hold for level *A_m_*, that (C3) is a new condition on the Cross operator, and that condition (C4) on Sel operator is different from (M3).

### 3.3 Analysis of selection mechanisms

We show how to parameterise the following selection mechanisms such that condition (M3) of Corollary 5 and (C4) of Corollary 6 are satisfied. In *k-tournament selection, k* individuals are sampled uniformly at random with replacement from the population, and the fittest of these individuals is returned. In (*μ*, λ)-*selection*, parents are sampled uniformly at random among the fittest *μ* individuals in the population. A function 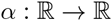 is a ranking function [28] if *α*(*x*) ≥ 0 for all *x* ∈ [0, 1], and 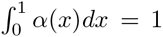. In ranking selection with ranking function *α*, the probability of selecting individuals ranked γ or better is 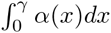. In *linear ranking selection* parametrised by *η* ∈ (1, 2], the ranking function is α(γ): = η(1 − 2γ) + 2γ. We define *exponential ranking selection* parametrised by *η* > 0 with α(γ): = *ηe*^η(1 − γ)^/(*e^n^* − 1).

#### Lemma 7.

Assuming that (*A*_1_,…, *A_m_*) is a partition of χ with (*A*_1_,…, *A*_*m* − 1_) being an f-based partition of χ \ *A_m_, for any constants δ*′ > 0, *p*_0_ ∈ (0, 1), ε ∈ (0, 1), and for any non-negative parameter *p_c_* = 1 − Ω(1), there exists a constant γ_0_ ∈ (0, 1) such that all the following selection mechanisms

1. k-tournament selection,
2. (μ, λ)-selection,
3. linear ranking selection,
4. exponential ranking selection

with their parameters *k*, γ/μ and *η* being set to no less than 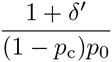 satisfy (M3), i. e. 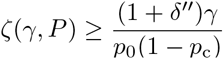 for any γ ∈ (0, γ_0_], any *P* ∈ (*χ*\*A_m_*)^λ^ and some constant δ″ > 0.

*Proof*. Since (M3) only concerns with the restricted subspace *χ* \ *A_m_* we only need to focus on this subspace, and because the partition is *f*-based on it, due to (16) it suffices to prove the results for *β* function instead of ζ function.

The results for *k*-tournament, (*μ*,*λ*)-selection and linear ranking follow by applying Lemma 13 in [16] (with its *p*_0_ being set as our *p*_0_(1 − *p*_c_)). For exponential ranking, we first remark the following lower bound,

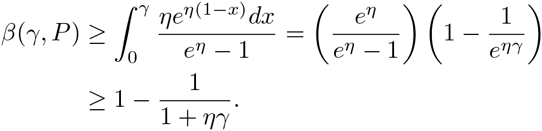

Then the rest of the proof is similar to *k*-tournament with *η* in place of *k*.

#### Lemma 8.

Assuming that (*A*_1_,…, *A_m_*) is a partition of χ with (*A*_1_,…, *A*_*m* − 1_) being an f-based partition of χ \ *A_m_*, for any constants δ′ > 0, *p*_0_ ∈ (0, 1) and ε ∈ (0, 1), there exists a constant γ_0_ ∈ (0, 1) such that the following selection mechanisms

1. *k*-tournament selection with *k* ≥ 4(1 + *δ*′)/(ε*p*_0_),
2. (*μ*, *λ*)-selection with λ/μ ≥ (1 + *δ*′)/(ε*p*_0_), *and*
3. exponential ranking selection with η ≥ 4(1 + *δ*′)/(ε*p*_0_)

satisfy (*C*4), i. e. 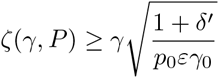 for any γ ∈ (0, γ_0_] and any *P* ∈ (*χ* / *A_m_*)^γ^.

*Proof*. Similar to the proof of Lemma 7, we only focus on the subspace *χ*\*A_m_* where the partition is *f*-based, and based on (16) we consider *β* function instead of ζ function.

Define ε′: = *εp*_0_.

1. Consider *k*-tournament selection and let γ ∈ (0, γ_0_]. By the definition of *f*-based partition, to select an individual from the same level as the γ-ranked individual or higher it is sufficient that the randomly sampled tournament contains at least one individual with rank γ or higher. Hence,

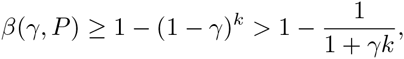

because 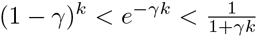. So for *k* ≥ 4(1 + *δ*′)/ε′,

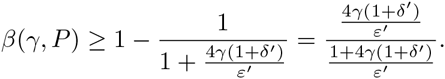

If γ_0_: = ε′/(4(1 + *δ*′)), then for all γ ∈ (0, γ_0_] it holds that 4(1 + γ′)/ε′ ≤ 1/γ and

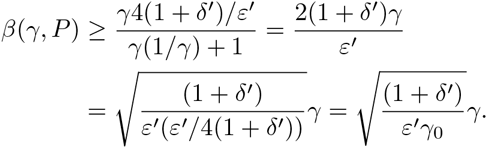

2. In (*μ*, γ)-selection, again by *f*-based property of the partition, we have *β*(γ, *P*) = λγ/*μ* if γ*δ* ≤ *μ*, and *β*(γ, *P*) = 1 otherwise. It suffices to pick γ_0_: = *μ*/λ so that with λ/*μ* ≥ (1 + *δ*′)/ε′, for all γ ∈ (0, γ_0_]. Then

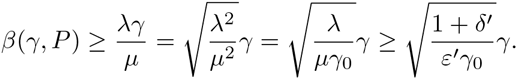

3. Similar to the proof of Lemma 7, we remark that 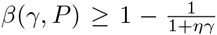, thus the rest of the proof is similar to *k*-tournament selection.

## 4 Applications to Genetic Algorithms for Different Problems

This section applies the results from the previous section to derive bounds on the expected runtime of GAs for optimising pseudo-Boolean functions and solving combinatorial Optimization problems.

In what follows, by bitwise mutation operator we mean an operator that given a bitstring *x*, computes a bitstring *y*, where independently of other bits, each bit *y_i_* is set to 1 − *x_i_* with probability *p_m_* and with probability 1 − *p_m_* it is set equal to *x_i_*. The tunable parameter *p_m_* is called a *mutation rate*.

### 4.1 Optimisation of pseudo-Boolean functions

In this subsection, we consider the expected runtime of non-elitist GAs in Algorithm 2 on the following functions,

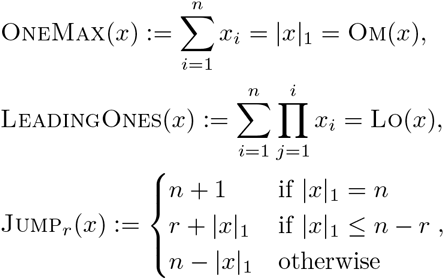

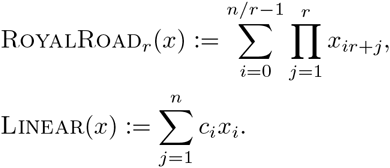

Note that our results on these functions also hold for their generalised classes, i. e. the meaning of 0-bit and 1-bit in each position can be exchanged, and/or *x* is rearranged according to a fixed permutation before each evaluation. For Linear, w. o. l. g. we can assume *c*_1_ ≥ *c*_2_ ≥ *c_n_* > 0 [16]. We also consider the class of *ℓ*-Unimodal functions, for which each function has exactly *ℓ* distinctive fitness values *f*_1_ < *f*_2_… < *f_ℓ_*, and each bitstring *x* of the search space is either optimal or it has a Hamming-neighbour *y* with a better fitness, i. e. *f*(*y*) > *f*(*x*).

For a moderate use of crossover, i. e. *p*_c_ = 1 − Ω(1), Corollary 5 is applicable and provides upper bounds on the expected runtime for all these functions and classes.

#### Theorem 9

The expected runtime of the GA in Algorithm 2, with *p_c_* = 1 − Ω(1) using any crossover operator, a bitwise mutation with mutation rate χ/n for any fixed constant χ > 0 and one of the selection mechanisms: k-tournament selection, (*μ*, λ)-selection, linear or exponential ranking selection, with their parameters *k*, λ/μ and η being set to no less than (1 + *δ*)*e^χ^*/(1 − *p_c_*) where δ > 0 being any constant, is

- 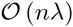 on OneMax if λ ≥ *c* ln *n*,
- 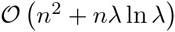 on LeadingOnes if λ ≥ *c* ln *n*,
- 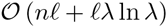 on ℓ-Unimodal if λ ≥ *c* ln(*ℓn*),
- 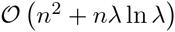 on Linear if λ ≥ *c* ln *n*,
- 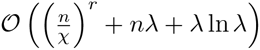 on Jump_*r*_ if λ ≥ cr ln *n*,
- 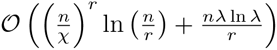 on RoyalRoad_*r* ≥ 2_ if λ ≥ *cr* ln *n*,

for some sufficiently large constant *c*.

*Proof*. We apply Corollary 5 with the canonical partition *A_j_*: = {*j* | *f*(*x*) = *j*} for all functions ^1^, except for Linear, the fitness-based partition [16]:

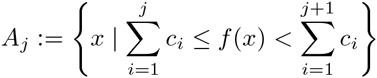

for *j* ∈ {0} ∪ [*n* − 1] and *A_n_*: = {1^n^}, is used.

The choices of *s_j_* and *s*_*_ to satisfy (M1) are the following.

- For OneMax, we set

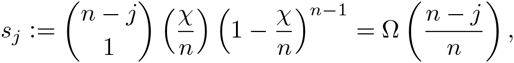

i. e. the probability of flipping a 0-bit while keeping all the other bits unchanged, and 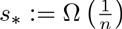.
- For LeadingOnes, *ℓ*-Unimodal and Linear, we set

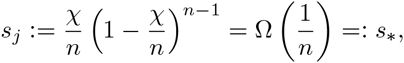

i. e. the probability of flipping a specific bit to create a Hamming neighbour solution with better fitness while keeping all the other bits unchanged. In *ℓ*-Unimodal, the bit to flip must exist by the definition of the function. In LeadingOnes, the 0-bit at position *j* + 1 should be flipped. For Linear, the partition satisfies that among the first *j* + 1 bits there must be at least a 0-bit, thus it suffices to flip the left most 0-bit will produce a search point at a higher level.
- For Jump_*r*_, as similar to OneMax for *j* ∈ [*n* − 1] we use

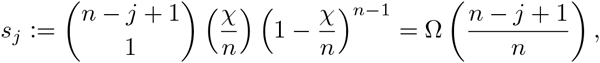

but 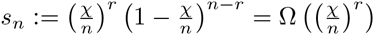, i. e. the probability of flipping the *r* remained 0-bits, so 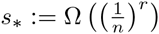.
- For RoyalRoad_*r*_, we use

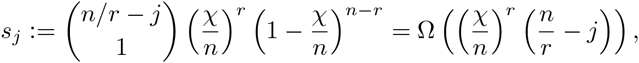
 i. e. the probability ofipping an entire unsolved block of length *r* (in the worst case) while keeping the other bits unchanged, and 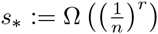.

It follows from Lemma 19 that the probability of not flipping any bit position by mutation is 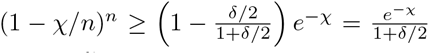 for *n* sufficiently large, thus choosing 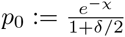 satisfies (M2).

We now look at (M3). In *k*-tournament selection, we have

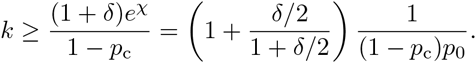

Hence, it follows from Lemma 7 that (M3) is satisfied with constant 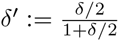. The same conclusion can be drawn for the other three selection mechanisms. In (M4), since γ_0_ and *δ*′ are constants, there should exist a constant *c* > 0 for each function such that the condition is satisfied given the minimum requirement on population size related to *c*.

Since all conditions are satisfied, Corollary 5 gives the desired result for each function. For OneMax and Jump_*r*_, optimisation time can be saved at early levels, i. e. *s_j_* is not small at the beginning, thus the evaluation of the sum 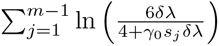 has to be precise:

- For OneMax, simplifying each term by ln 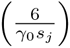 gives

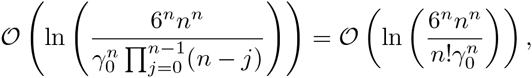

and by Stirling’s approximation 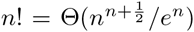, this is no more than 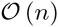. The expected runtime is then 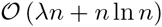. Since we already require λ = Ω(ln *n*), this can be written shortly as 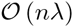.
- For Jump_*r*_, we use the simplication ln 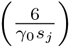 for therst *m* − 2 terms of the sum, and ln(3λ/2) for the last term, so this gives

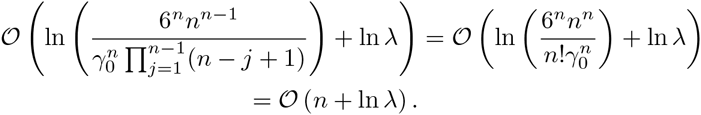 The expected runtime is then 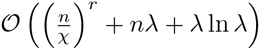.

For the other functions, *s_j_* is already small at early levels, thus there is no benefit of considering the gradual sum of ln. Hence, the simpli cation 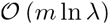 for the sum 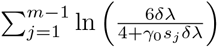 gives the corresponding results.

In the case of regular use of crossover, i. e. *p*_c_ = 1, the relationship between the crossover operator and the structure of the search space becomes non-negligible. In the following, we consider a general *mask-based* crossover as follows. Given two parent genotypes u; v; the operator consists in first choosing (deterministically or randomly) a binary string 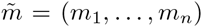 to produce two o spring vectors *x*′, *x*″ as

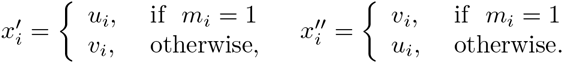

Then one element of {*x*′, *x*″} chosen uniformly at random is returned. For example, the uniform crossover is a mask-based crossover for which 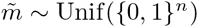, and a *k*-point crossover is a mask-based crossover for which

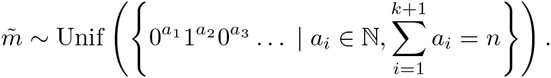

The following lemma shows that all mask-based crossover operators satisfy (C3) with ε = 1/2 for Om and Lo functions.

#### Lemma 10.

If *x* ~ *p*_xor_(*u, v*), where *p_xor_* is a mask-based crossover, then:

1. 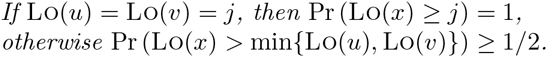
2. 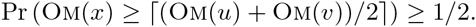

*Proof*. 1) When Lo(*u*) = Lo(*v*) = *j*, in mask-based crossover operators, the two bitstrings *x*′, *x*′ have *j* leading ones. So does the returned bitstring, i. e. with probability 1.

If Lo(*u*) = Lo(*v*), we can assume w. o. l. g. that Lo(*v*) = *j* and Lo(*u*) > Lo(*v*). Then *v* has a 0 while *u* has a 1 at position *j* + 1. So, one of the bitstrings *x*′, *x*′′ in the mask-based crossover will inherit the 1 at that position and the other will inherit the 0. This implies that one of them has fitness at least *j* + 1 and with probability 1/2 it is returned as output.

2) Each bit of *u* and *v* is copied either to *x*′ or to *x*′′, therefore |*x*^′^|_1_ + |*u*|_1_ + |*v*|_1_, which means that max 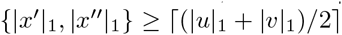. The output is chosen with probabilities 1/2 to be copied either from *x*′ or *x*′′, and the result follows.

#### Theorem 11.

Assume that the GA in Algorithm 2 with p_c_ = 1 uses any mask-based crossover operator, a bitwise mutation with mutation rate =n for any fixed constant > 0, and one of the following selection mechanisms:

- *k*-tournament selection with *k* ≥ 8(1 + δ)e^x^,
- (*μ*,*λ*)-selection with 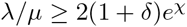, or
- exponential ranking selection with 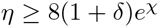,

for any constant *>* > 0. Then there exists a constant *c* > 0, such that the expected runtime of the GA is

- 𝒪 (*n*λ) on ONEMAX if *λ* ≥ cln *n*,
- 𝒪 (*n*^2^ + *n*λ ln λ) on LEADINGONES if *λ* ≥ cln *n*.

Proof. We apply Corollary 6 this time, but again using the canonical partition of the search space for both functions. We also assume that n is large enough so that by Lemma 19 the probability of not flipping any bit by mutation is 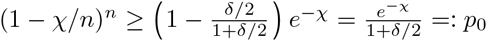, and so (C2) is satisfied with this choice of p_0_. In addition, we use the same upgrades probabilities *s_j_* and their smallest value s for each of the two functions as in the proof of Theorem 9 to satisfy (C1).

It follows from Lemma 10 that (C3) is satisfied for constant ε_1_: = 1/2. We now look at condition (C4). For k-tournament, we get 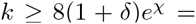 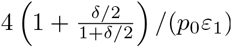. So condition (C4) is satisfied with constant 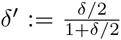 for k-tournament by Lemma 8. The same reasoning can be applied so that (C4) is also satisfied for the other selection mechanisms.

Since *δ*′ and γ_0_ are constants, thus condition (C5) is satisfied given λ ≥ *c* ln *n* and for some constant *c*. Since all conditions are satisfied, the result follows from Corollary 6.

Note that the upper bounds in Theorem 11 match the upper bounds of Theorem 9. The latter is a generalisation of the results in [16] which were limited to EAs without crossover.

In the next sections, we further demonstrate the generality of Theorem 1 through Corollary 5 by deriving bounds on the expected runtime of GAs with p_c_ = 1 Ω (1) to optimise or to approximate the optimal solutions of combinatorial optimisation problems. We start with a simple problem of sorting n elements from a totally ordered set.

### 4.2 Optimisation on permutation space

Given n distinct elements from a totally ordered set, we consider the problem of ordering them so that some measure of sortedness is maximised. Several measures were considered by [52] in the context of analysing the (1+1) EA. One of those is Inv() which is defined to be the number of pairs (*i*; *j*) such that 1 ≤ *i* < ≤ *j* n, π(*i*) < π(*j*) (i.e. pairs in correct order). We show that with the method introduced in this paper, i. e. Corollary 5 analysing GAs on Sorting problem with Inv measure, denoted by Sorting_Inv_, is not much harder than analysing the (1+1) EA.

For the mutation we use the Exchange(7π) operator [52], which consecutively applies *N* pairwise exchanges between uniformly selected pairs of indices, where *N* is a random number drawn from a Poisson distribution with parameter 1.

#### Theorem 12.

If the GA in Algorithm 2 with *p_c_* = 1 Ω(1) uses any crossover operator, the Exchange mutation operator, one of the selection mechanisms: k-tournament selection, (;)-selection, and linear or exponential ranking selection, with their parameters *k*, *λ*/*μ* and being set to no less than (1+δ)e/(1 − *p_c_*), then there exists a constant *c* > 0 such that if the population size is λ ≥*c* ln *n*, the expected time to obtain the optimum of Sorting_Inv_ is O (*n*^2^λ).

Proof. Define 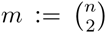. We apply Corollary 5 with the canonical partition, 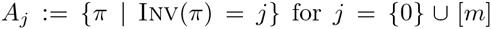. The probability that mutation exchanges 0 pairs is 1/e. Hence, condition (M2) is satisfied for *p*_0_: = 1/*e*.

To show that (M1) is satisfied, we first define 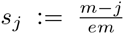 for each *j* ∈ {0} ⋃ [*m* - 1]. Since *x* ∈ *A_j_*, then the probability that the exchange operator exchanges exactly one pair is 1/e, and the probability that this pair is incorrectly ordered in *x*, is (*m - j*)/*m*. Thus, (M1) is satisfied with the defined *s_j_*.

In (M3), for k-tournament we have that 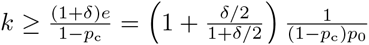 thus the condition is satisfied for constant 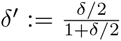 and some constant γ_0_ ∈ (0,1) by Lemma 7. The same conclusion can be drawn be the other selection mechanisms. Finally, since γ_0_; δ^'^ are constants, there exists a constant c > 0 such that (M4) is satisfied for any *c* ≥ ln(*n*).

It therefore follows by Corollary 5 that the expected runtime of the GA on Sorting_Inv_ is O n^2^λ, i. e. this is similar to 𝒪neMax except that we have *m* = 𝒪(*n*^2^) levels.

### 4.3 Search for Local Optima

A great interest in the area of combinatorial optimisation is to nd approxi-mate solutions to NP-hard problems, because exact solutions for such problems are unlikely be computable in polynomial time under the so-called P6=NP hy-pothesis. In the case of maximisation problems, a feasible solution is called a ρ-approximate solution if its objective function value is at least times the optimum for some ρ ∈ (0;1]. Local search is one method among others to approximate solutions for combinatorial optimisation problems through finding local optima (a formal definition is given below). For a number of well-known problems it was shown [1] that any local optimum is guaranteed to be a ρ- approximate solution with a constant.

Suppose that a neighbourhood *N (x)* ⊆ χ is defined for every *x* ∈ χ. The mapping *N*: χ → 2^χ^ is called the neighbourhood mapping and all elements of *N (x)* are called neighbours of x. For example, a frequently used neighbourhood mapping in the case of binary search space χ = {0,1}^n^ is defined by the Hamming distance H(.,.) and a radius r as

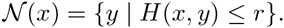

If f(y) ≤ f(x) holds for all neighbours y of *x* ∈ χ, then *x* is called a local optimum w. r. t. N. The set of all local optima is denoted by 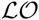 (note that global optima also belong to 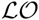).

A local search method starts from some initial solution y_0_. Each iteration of the algorithm consists in moving from the current solution to a new solution in its neighbourhood, so that the value of the fitness function is increased. The way to choose an improving neighbour, if there are several of them, will not matter in this paper. The algorithm continues until a local optimum is reached. Let m be the number of different fitness values attained by solutions from 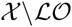 plus 1. Then starting from any point, the local search method finds a local optimum within at most m - 1 steps.

Alternatively, one can use GAs to solve the optimisation problem, and possi-bly nd local optima. The following result provides sets of sufficient conditions for a performance guaranteed GA to find local optima.

**Corollary 13.** *Given some positive constants pε_0_; _0_ and, define the following conditions:*

**(X1)** p_mut_(y | x) ≥ s for any *x* ∈ χ, y ∈ *N (x)*.
**(X2)** p_mut_(x | x) ≥ p_0_ for all *x* ∈ χ.
**(X3)** p_xor_ (f(x^'^) ≥ max{f(u); f(u), f (v))≥ ε_0_ for any u, v ∈ χ.
**(X4.1)** the non-elitist GA in Algorithm 2 is set with p_c_ = 1 - Ω(1), and it uses one of the following selection mechanisms:
  - *k-tournament selection with* 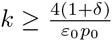,
  - (*μ*, λ)-selection with 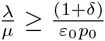,
  - *exponential ranking selection with* 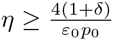.
**(X4.2)** the non-elitist GA is set with p_c_ = 1 - Ω(1), and it uses one of the following selection mechanisms: k-tournament selection, (μ,λ)-selection, linear or exponential ranking selection, with their parameters k, λ/μ and η being set to no less than 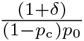.

*If (X1-3) and (X4.1) hold, or exclusively (X1-2) and (X4.2) hold, then there exists a constant c, such that for* 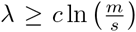, *a local optimum is reached for the first time after* 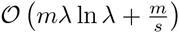 fitness evaluations in the expectation.

Condition (X4.1) or (X4.2) characterises the setting of selection mechanisms, while (X1-3) bear the properties of the variation operators over the neighbourhood structure *N*. Particularly, (X1) assumes a lower bound s on the probability that the mutation operator transforms an input solution into a specific neighbour. To illustrate this condition, we note that in most of the local search algorithms the neighbourhood *N* (x) may be enumerated in polynomial time of the problem input size. For such neighbourhood mappings, a mutation operator that generates the uniform distribution over *N* (x) will select any given point in *N* (x) with probability at least s; so that 1=s is polynomially bounded in the problem input size.

If crossover is frequently used, i. e. *p_c_* = 1, we also need to satisfy condition (X3) on the the crossover operator. It requires that the fitness of solution *x* on the output of crossover is not less than the fitness of parents with probability at least ε_0_. Note that such a requirement is satisfied with ε_0_ = 1 for the optimized crossover operators, where the o spring is computed as a solution to the *optimal recombination problem* (see e.g [23]). This supplementary problem is known to be polynomially solvable for Maximum Clique [4], Set Packing, Set Partition and some other NP-hard problems [23].

When a set of conditions, i. e. depending on whenever *p_c_* = 1 or p_c_ = 1 (1), is satisfied and given a sufficiently large population w. r. t. to *m* and *s*, an upper bound on the expected number of fitness evaluations that GA performs to find a local optimum is guaranteed. The proof directly follows from Corollaries 6 and 5 of Theorem 1.

*Proof of Corolary 13*. We use the following partition

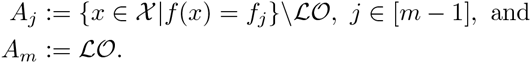

We note that (A_1_,…, A_m 1_) is a fitness-based partition of 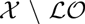. Thus, applications of Corollary 5 and Lemma 7 for the set of conditions (X1-2) and (X4.2), or alternatively, Corollary 6 and Lemma 8 for the set of conditions (X1-3) and (X4.1), yield the required result.

A similar result for GAs with very high selection pressure was obtained in [25]. In particular, the result from [25] implies that a GA with tournament selection or (;)-selection, given certain settings of parameters, reaches a local optimum after O (m ln(m)/s) fitness evaluations in expectation. The upper bound from Corollary 13 has an advantage to the bound from [25] if 1/s is at least linear in *m*.

The effect of the corollary can be seen in the following example setting. Let us consider the binary search space {0,1}^n^ with Hamming neighbourhood mapping of a constant radius *r*, *a* fitness function *f* such that *m* ∈ poly(n), and assume that GA uses bitwise mutation operator and p_c_ = 1 - χ(1). The bitwise mutation operator outputs a string *y*, given a string *x*, with probability p^H^_m_^(x,y)^(1 - p_m_)^n-H(x,y)^. Note that probability p^j^_m_(1-*p_m_*)^n-j^, as a function of p_m_ attains its maximum at *p_m_* = *j/n*. It is easy to show (see e.g. [25]) that for any *x* ∈ χ and *y* ∈ *N* (x), the bitwise mutation operator with *p_m_* = *r/n* satisfies the condition *p_mut_(y | x) = O (1/n^r^*). Besides that, for a sufficiently large *n* and any *x ∈ χ* holds *p_mut_(x | x)*≤ e ^-r^/2 = Ω(1). Therefore, Corollary 13 implies that a GA with the above mentioned operators given appropriate settings of parameters; *p_m_* and *p_c_*, first visits a local optimum w. r. t. a Hamming neighbourhood of constant radius after a polynomially bounded number of fitness evaluations in expectation.

To give concrete examples, we consider the following unconstrained (and unweighted) problems:

- Max-SAT: given a CNF formula in n logical variables which is represented by *m'* clauses c_1_,…, c*_m_*' and each clause is a disjunction of logical variables or their negations, it is required to find an assignment of the variables so that the number of satisfied clauses is maximised.
- Max-CUT: given an undirected graph G = (V; E), it is required to find a partition of V into two sets (S, V \ S), so that the number of crossing edges, i. e. 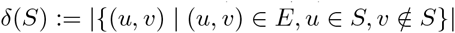, is maximised.

Both problems are NP-hard, and their solutions can be naturally represented by bitstrings. Particularly, any local optimum w. r. t. the neighbourhood defined by Hamming distance 1 has at least half the optimal fitness [1]. Better approximation ratios can be obtained with more sophisticated algorithms, e. g. 0:79-approximation for Max-SAT [2] based on a time-consuming semi-definite programming relaxation. The local search algorithm with the above neighbourhood however has an advantage of low time complexity, e. g. it only makes O (*nm^0^*) tentative solutions for Max-SAT. Corollary 13 translates such a result into relatively low runtime bound for GAs.

**Theorem 14.** Suppose the GA in Algorithm 2 is applied to Max-SAT or to Max-CUT using a bitwise mutation with p_m_ = =n, where > 0 is a constant, a crossover with p_c_ = 1 (1) and one of the selection mechanisms: k-tournament selection, (μλ)-selection, linear or exponential ranking selection, with their parameters k, λ/μ and being set to no less than 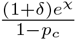, where δ > 0 is any constant. Then there exists a constant c, such that for λ ≤ cln(nm^ȧ^), a1/2-approximate solution is reached for the first time after 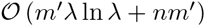 fitness evaluations in expectation for Max-SAT, and after 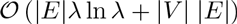 for Max-CUT.

The proof is analogous to the analysis of `-Unimodal function in Theorem 9, combined with Corollary 13 where m is no more than m^'^ + 1 for Max-SAT and no more than |E| + 1 for Max-CUT.

## 5 Estimation of Distribution Algorithms

As mentioned in the introduction, there are few rigorous runtime results for UMDA and other estimation of distribution algorithms (EDAs). The analytic techniques used in previous analyses of EDAs were often complex, e. g. relying on the machinery of Markov chain theory. Surprisingly, even apparently simple problems, such as the expected runtime of UMDA on OneMax, were until recently open.

Algorithm 1 matches closely the typical behaviour of estimation of distribution algorithms: given a current distribution over the search space, sample a finite number of search points, and update the probability distribution. We demonstrate the ease at which the expected runtime of UMDA with margins and truncation selection on the OneMax function can be obtained using the level-based theorem without making any simplifying assumptions about the optimisation process.

### 5.1 Algorithm

If *P ∈ X*^λ^ is a population of λ solutions, let *P (k,i)* denote the value in the *i*-th bit position of the *k*-th solution in *P*. The Univariate Marginal Distribution Algorithm (UMDA) with (μ,λ)-truncation selection is defined in Algorithm 4.

The algorithm has three parameters, the parent population size, the μ - spring population size λ, and a parameter *m^'^ μ* controlling the size of the margins. It is necessary to set *m^'^ > 0* to prevent a premature convergence, e. g. without this margin *p_t_(i)* can go to a non-optimal fixation, this prevents further exploration and causes an infinite runtime. Based on insights about optimal mutation rates in the (1+1) EA, we will use the parameter setting *m^'^ = =n* in the rest of this section.

It is immediately clear that the UMDA in Algorithm 4 is a special case of Algorithm 1 scheme. The probability distribution *D(P_t_)* of *y* is computed in steps 6-7, and is defined for any search point *x* ∈ {0,1}*^n^* by

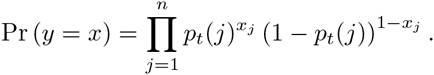

#### Algorithm 4

UMDA

1: Initialise the vector p_0_: = (1/2,…, 1/2).

2: for t = 1; 2; 3;… do

3: for *x* = 1 to do

4: Sample the x-th individual P_t_(x,.) according to

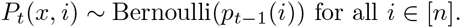

5: end for

6: Sort the population P_t_ according to f.

7: Calculate a new vector p_t_ from P_t_ according to

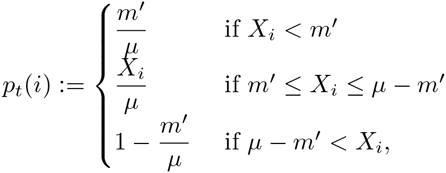

for all i ∈ [n], where X_i_: =_k=1_P_t_(k,i).

8: end for

Note that the sampling from vector (*p_i_)_i∈[n]_* in UMDA is analogous to a population-wise crossover, i. e. the bit sampling at position i with a non-marginal probability (line 4 in Algorithm 4) is equivalent to picking uniformly at random a bit from the bits at position *i* of the μ selected individuals of the previous generation. In some other randomised search heuristics such as ant colony optimisation (ACO) and compact genetic algorithms (cGA), the sampling distribution *D_t_* does not only depend on the current population, but also on additional information, such as pheromone values. The level-based theorem does not apply to such algorithms.

It is well-known that the (1+1) EA solves OneMax problem in expected time *͈(n ln n)*, and this is optimal for the class of unary, unbiased black-box algorithms. Surprisingly, no previous runtime analysis of UMDA seems available for OneMax. We demonstrate that the expected runtime can be obtained relatively easy with our methods. To obtain lower bounds on the tail of the level-distribution, we make use of the Feige inequality [26] (or see Lemma 20 in the appendix).

**Theorem 15.** *Given any positive constants* δ ∈ (0,1), and 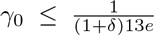 *the UMDA with o spring population size* with *b ln(n)≤ λ≤n/γ*_0_ *for some constant b* > 0, parent population size μ = γ_0_λ and margins *m*^'^ = μ/*n*, has expected optimisation time *O (nλ ln)* on OneMax.

*Proof*. Step 1: We use the canonical partition into m = n + 1 levels, where level j ∈ [m] is defined by

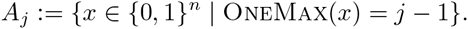

We use the parameter *m_'_*: = μ/λ and let *Y* be the number of one-bits in the sampled solution.

The choice *m^'^* = μ/*n* and μ ≤ n implies that the margins for *p_t_(i)* are simplified to 1/*n* and 1 - 1/*n*, and that these margins are only used when the bit values at position *i* of the selected individuals are identical. We categorise the probabilities *p_t_(i)* into three groups: those at the upper margin 1 - 1/*n*, those at the lower margin 1/*n*, and intermediary values in the closed interval [1/μ, 1 - 1/μ]. Due to linearity of the fitness function, the components of *p_t_* can be rearranged without changing the distribution of *Y*. We assume w.l.o.g. a rearrangement so that there exists integers *k,l* ≥ 0 satisfying

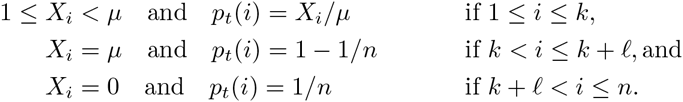

By these assumptions, it follows that

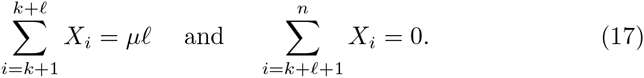

In the following, we define *Y_i,k_* to be the number of sampled one-bits due to (*p_t_(i)*,…, *p_t_(k)*) in the rearranged *p_t_*.

For any population *P_t_* and any γ ∈ [0; _0_], let *j* ∈ [*n*] be any integer such that |*P_t_* ⋂ A _≥j_|≥ γ_0_ λ=μ and |*P_t_* ⋂ A _≥j+1_|≥ γλ. This implies that among the *μ* fittest individuals in the current population, there are at least γλ individuals with at least *j* one-bits, and the remaining among the fittest individuals have at least *j* - 1 one-bits. Hence, the total number of one-bits among the fittest individuals must satisfy

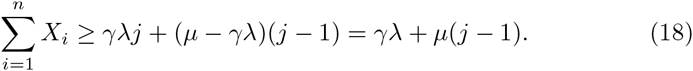

Combining Eqs. (17) and (18), when *k* ≥ 1, we get

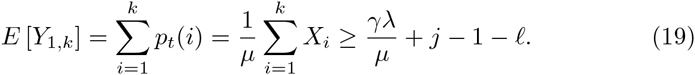

Step 2: We first verify condition (G2), i. e. checking if Pr (Y ≥ j) ≥ (1 +δ)γ for any level *j* defined like above with > > 0. We distinguish between two cases, either *k* = 0 or *k* ≥ 1.

Case 1: If *k* ≥ 1, then Eq. (19) and Lemma 20 give

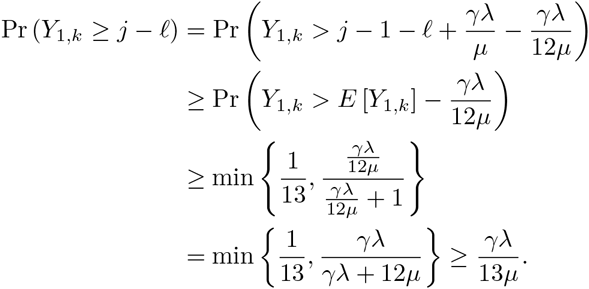

The probability of sampling an individual with at least *j* one-bits in the next generation is therefore lower-bounded by

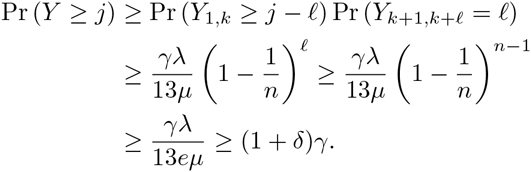

Case 2: If *k* = 0, then all the μ best individuals in the population must be identical. By assumption, there are γ,λ ≥ 1 individuals with at least *j* 1-bits, hence all the best individuals must have at least *j* 1-bits. In this case, there are *l* ≥ *j* probabilities at the upper margin, and we get

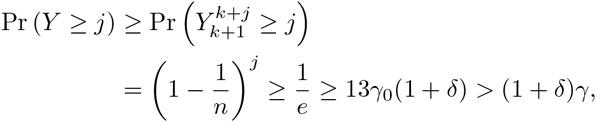

and condition (G2) is therefore satisfied also in this case.

Step 3: We now consider condition (G1) for any *j* defined with = 0. Again we check the two cases *k* = 0, and *k* ≥ 1.

Case 1: If *k* = 0, then with our assumption, the *l* ≥ *j* - 1 first probabilities are at the upper margin 1 - 1/*n*, and the last *n - l* ≤ *n-j*+1 probabilities are at the lower margin 1/*n*. In order to obtain a search point with at least *j* one-bits, it is sufficient to sample exactly *l*≥ *j* - 1 one-bits in the first *l* positions and exactly one 1-bit in the last *n - l* ≤ *n - j* + 1 positions. Hence,

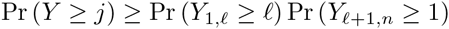

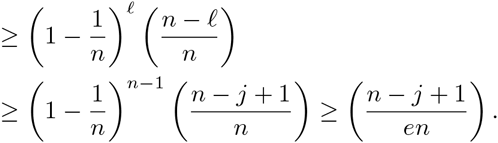

Case 2: When *k* ≥ 1, we note from Eq. (19) that

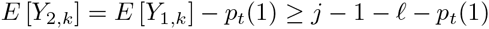

Again, by Lemma 20 we get

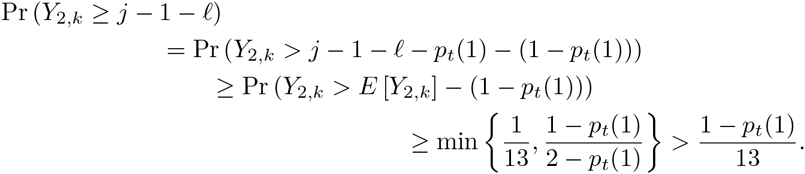

The probability of sampling an individual with at least *j* one-bits in this con-guration is bounded from below as

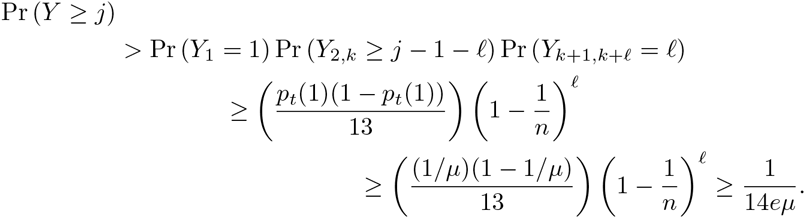

The last inequality holds for μ ≥ 14, which in turn only requires n to be larger than some constant. Hence, combining the cases *k* = 0 and *k* > 0, for all *j* ∈ [n] we get

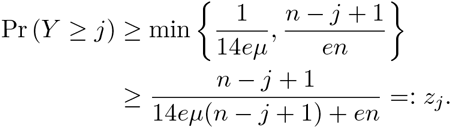

Clearly, there exists a *z_∗_* with 1/*z_∗_* ∈ poly(*n*) such that Pr (*Y ≥ j*) *z* for all *j* ∈ [*n*] and condition (G1) is satisfied.

Step 4: We consider condition (G3) regarding the population size. The parameters δ and _0_ = = are constants with respect to n, therefore the variables a; ε and c in condition (G3) are also constants, and 1/*z_∗_* ∈ poly(*n*). Hence, there must exist a constant *b* > 0 such that condition (G3) is satisfied when λ ≥ blog(*n*):

Step 5: To conclude, the expected optimisation time is

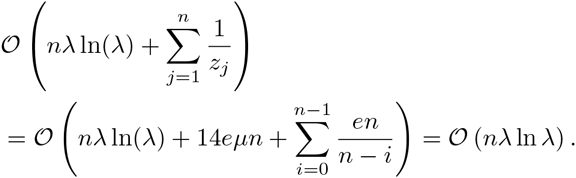

A similar analysis for LeadingOnes [15] yields an upper bound on the runtime of 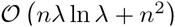 with o spring population size b ln(*n*) for some constant *b* > 0 without use of Feige’s inequality. The previous result [9] on LeadingOnes requires a larger population size and gives a longer runtime bound.

Table 1 summarises the runtime bounds for the example applications of the tools presented in this paper and the above mentioned result for UMDA on LeadingOnes.

**Table 1.**
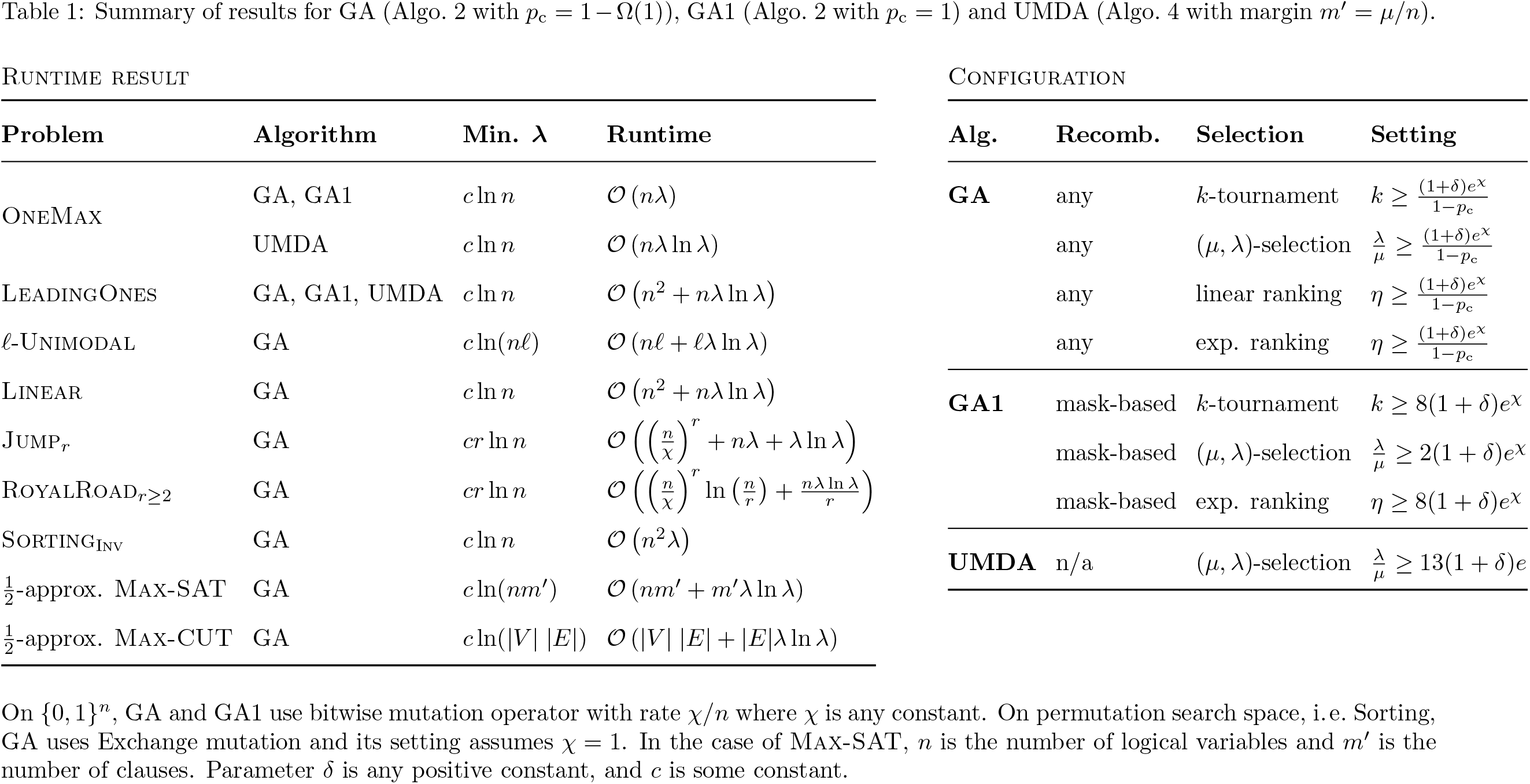
Summary of results for GA (Algo. 2 with *p*_c_ = 1 − Ω(1)), GA1 (Algo. 2 with *p*_c_ = 1) and UMDA (Algo. 4 with margin *m*′ = *μ*/*n*).

## 6 The level-based theorem is almost tight

How accurate are the time bounds provided by the level-based theorem? To answer this question, we first interpret the theorem as a universally quantified statement over the operators D satisfying the conditions of the theorem. More formally, given a choice of level-partitioning and set of parameters *z*_1_,…, *z*_m 1_;; _0_, which we collectively denote by Θ, the theorem can be ex-pressed on the form

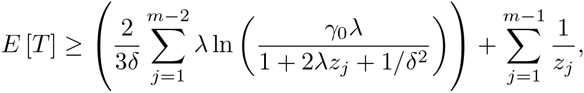

where 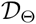 is the set of operators *D* in Algorithm 1 that satisfy the conditions of the level-based theorem with parameterisation Θ, *E* [*T_D_*] is the expected running time of Algorithm 1 with a given operator *D*, and t is the upper time bound provided by the level-based theorem which depends on the parameterisation Θ.

In order to obtain an accurate bound for a specific operator *D*, for example the (μ,λ) *EA* applied to the OneMax function, it is necessary to choose a parameterisation that reffects this process as tightly as possible. If the bounds on the "upgrade" probabilities *z_j_* for the (μ,λ) EA are too small, then the class 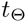 includes other processes which are slower than the (μ,λ) EA, and the corresponding bound t cannot be accurate. Hence, the theorem is limited by the accuracy at which one can describe the process by some class 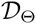. One could imagine a more accurate variant of the theorem requiring more precise, and possibly harder to obtain, information about the process, such as the variance of *D(P_t_)*.

Assuming a fixed parameterisation Θ, it is possible to make a precise statement about the tightness of the upper bound t_Θ_. Theorem 16 stated below is an existential statement on the form

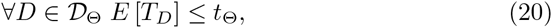

where the lower bound 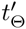 is close to the upper bound 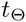. Hence, given the information the theorem has about the process through the parameterisation Θ, the runtime bound is close to optimal. More information about the process would be required to obtain a more accurate bound on the runtime.

In some concrete cases, one can prove that the level-based theorem is close to optimal using parallel black-box complexity theory [3, 22]. From Corollary 5 with *p_c_* = 0, which specialises the level-based theorem to algorithms with unary mutation operators, one can obtain the bounds 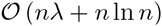 for OneMax, and 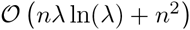 for LeadingOnes for appropriately parameterised EAs. These bounds are within a O (lnλ)-factor of the lower bounds that hold for any parallel unbiased black-box algorithm [3]. For population sizes 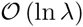 and 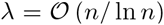, the resulting 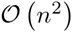 bound on LeadingOnes is asymptotically tight, because it matches the lower bound that holds for all black-box algorithms with unary unbiased variation operators [40].

**Theorem 16.** Given any partition of *x* into m non-empty subsets (A_1_,…, A_m_), for any z_1_,…, z_m-1_ δ γ_0_ ∈ (0,1) where 1 ≥ _0_(1 + δ) ≥ z_j_ for all *j* ∈ [m - 1], and 2 ℕ, there exists a mapping *D* which satisfies conditions (G1), (G2), and (G3), of Theorem 1, such that Algorithm 1 with mapping *D* has expected hitting time

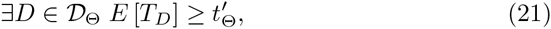

where 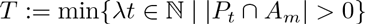.

Proof of Theorem 16. We construct an operator *D* which leads to the claimed lower bound. Choose any sequence of search points (*x*_1_,…, *x*_m_) 2 *A*_1_ *A_m_*, and let the initial population of Algorithm 1 be *P_0_*: = (*x*_1_,…, *x*_1_), i.e., copies of the search point *x*_1_ belonging to the first level.

For any population *P* ∈ χ^λ^, let the current level be the largest *i* ∈ [m] such that |*P* ⋃ *A* _≥i_|≥ γ_0_λ. For any population *P* with current level *i* < *m*, define the operator *D* for all μ ∈ χ by

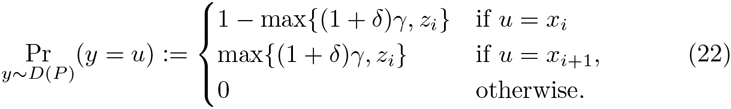

where:= (1/λ)P ⋃ A _≥i+1_| < γ_0_.

For all 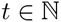, define

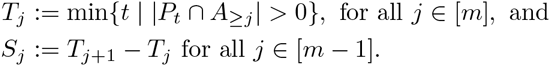

Then we have 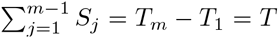 because T_1_ = 0. The random variable *S*_j_, for *j* ∈ [*m − 1*], describes the number of generations from the time the process has discovered the search point *x*_j_ until it has discovered the search point *x*_j+1_, and we call this phase *j*. We divide each phase *j* into two sub-phases. Let S_j_^1^ be the number of generations where

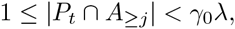

and call this the first sub-phase, and let 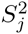 be the number of generations where

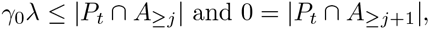

and call this the second sub-phase. The duration of the *j*-th phase is the sum 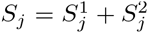. Remark that 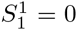 due to the choice of the initial population *P*_0_.

Note also that by the definition of operator *D*, as long as the process is in sub-phase 1 of phase *j*, the probability of generating the search point *x*_j+1_ is 0. Furthermore, the process never returns to sub-phase 1 once the process has entered sub-phase 2. To estimate the duration of sub-phase 1, we consider the stochastic process 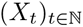 where 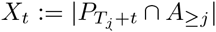, and a corresponding filtration 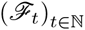 where 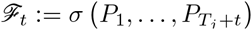.

During sub-phase 1 of phase *j* > 1, it holds that *x*_t+1_ ~ Bin(; p_t+1_), where 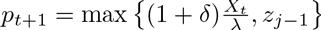.

To lower bound the expected duration of sub-phase 1, we apply drift analysis (Lemma 28) with respect to the process 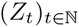 defined by Z_t_: = ln(λ*R_t_*) where *R_t_*: = max{*x*_t_; *y_j_*} and *y_j_*: = max{*z_j-1_*,1/δ^2^} > 1: Note that since *z_j_* < γ_0_ by assumption, and 1= ^2^ < γ_0_λ by condition (G3), it holds that *y_j_* < γ_0_λ. It is therefore clear that sub-phase 1 is only complete if

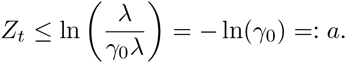

By Jensen’s inequality, the drift of this process can be bounded by

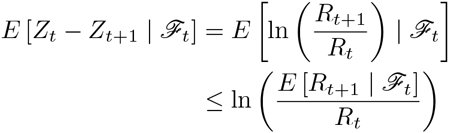

and by Lemma 29

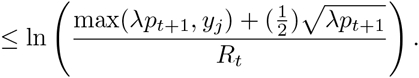

When *p*_t+1_ *y_j_*, we use that *R_t_* = max{*x*_t_; *y_j_*}≥ λ_pt+1_/(1 +δ) because p_t+1_ = max{*x*_t_(1 +δ)/λ z_j-1_}, so

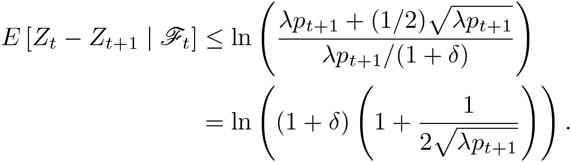

Since *p*_t+1_ ≥ *y*_j_ > 1, thus 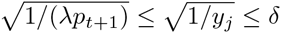 and

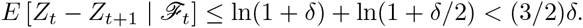

Otherwise, when *y*_j_ > *p*_t+1_, we use that *R*_t_ *y*_j_ and get

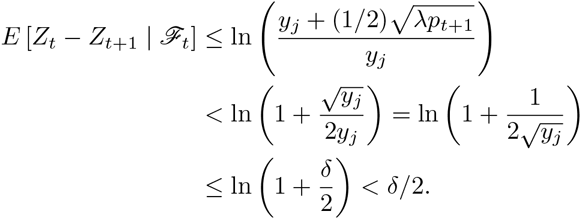

Hence, condition 1 in Lemma 28 can be satisfied with the parameter ε:= (3/2)δ. We therefore get the bound

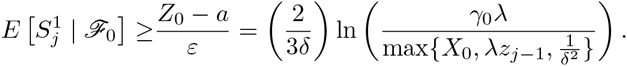

By the definition of the process, for 1 < *j* ≤ m; we have *x*_0_~(*Y* | ≥ *Y*1) where Y ~ Bin(λ, z_j-1_); i.e., *x*_0_ is binomially distributed random variable conditional on having value at least 1. and by the tower property of expectation,

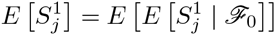

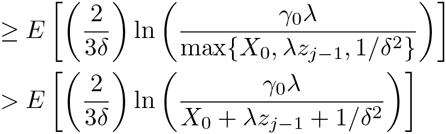

and since the function *f(x)* = ln(1/*x*) is convex, Jensen’s inequality and Lemma 30 give

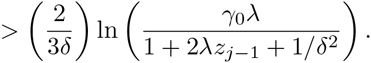

During sub-phase 2, it holds that

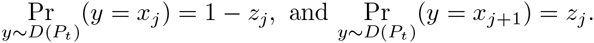

In each generation of sub-phase 2, the phase ends with probability *q_j_*: = 1 - (1 − *z_j_*)^λ^ < λz_j_, i.e., the probability that at least one individual is produced in *A* _≥j+1_. The duration of sub-phase 2 is therefore geometrically distributed with parameter *q_j_* and has expectation 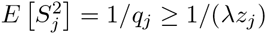.

Hence, we get

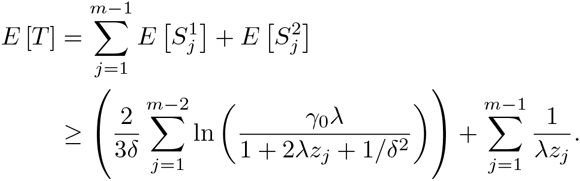

## 7 Conclusion

Time-complexity analysis of evolutionary algorithms (EAs) has advanced significantly over the last decade, starting from simplified settings such as variants of the (1+1) EA without a real population, crossover or other higher-arity operators. It has been unclear to what extent the time-complexity pro les of these simple EAs considered by theoreticians deviate from those of the more sophisticated, population-based EAs often preferred by practitioners. New techniques tailored to time-complexity of population-based algorithms are required.

This paper introduces a new technique that easily yields upper bounds on the expected runtime of complex, non-elitist search processes. The technique is first illustrated on Genetic Algorithms. We have shown that GAs optimise standard benchmark functions, as well as combinatorial optimisation problems, *e* ciently. As long as the population size is not overly large, the population does not incur an asymptotic slowdown on these functions compared to standard EAs that do not use populations. Thus, speedups can be achieved by parallellising fitness evaluations. Furthermore, consequent work indicate that non-elitist, population-based EAs have an advantage on more complex problems, including those with noisy [14], dynamic [13], and peaked [17] fitness landscapes.

As a side-effect of the analysis, the conditions of level-based theorem yield settings for algorithmic parameters, such as population size, mutation and crossover rates, selection pressure etc., that are sufficient to guarantee a given time-complexity bound. This opens up the possibility of theory-led design of EAs with guaranteed runtime, where the algorithm is designed to satisfy the conditions of the level-based theorem [11].

Further demonstrating the generality of the theorem, we also provide time-complexity results for the UMDA algorithm, an Estimation of Distribution Algorithm, for which there are few theoretical results. Finally, we show via lower bounds on the runtime of a concrete process, that given the information the theorem requires about the process, the upper bounds are close to tight, i.e., little improvement of the time bound in the theorem is possible.

## Acknowledgment

The research was supported by the European Union Seventh Framework Programme (FP7/2007-2013) under grant agreement no 618091 (SAGE) and Russian Foundation for Basic Research grants 15-01-00785 and 16-01-00740. Early ideas were discussed at Dagstuhl Seminars 13271 and 15211 “Theory of Evolutionary Algorithms”.

## Appendix A

The following results are known in the literature.

**Lemma 17** (Lemma 33 in [16]). For all *x* ≥ 0, *x* ln ≥(1 + *x*)≥ x(1 - *x*/2).

**Lemma 18** (Lemma 31 in [16]). For 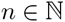 and *x* ≥ 0, we have 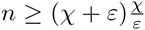.

**Lemma 19** (Lemma 3 in [17]). For any ε∈ (0,1) and *χ* > 0, if 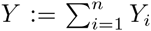 then 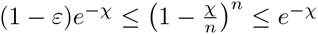.

**Lemma 20** (Corollary 3 in [15]). Let Y1,…, Yn be n independent random variables with support in [0,1] and finite expectations and 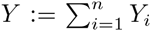. It holds for every *δ* > 0 that

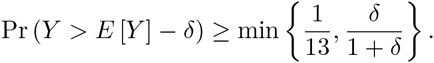

## Appendix B

The following lemmas are part of the proof of Theorem 1.

**Lemma 21.** The functions *g*_1_ and *g*_2_ defined below are level functions for any 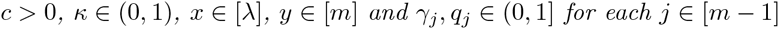.

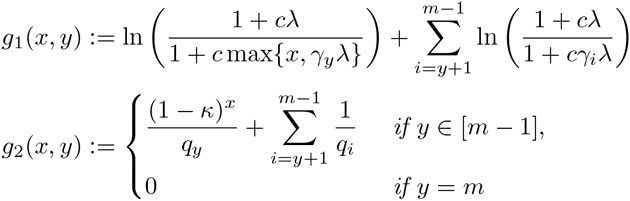

and *g*_1_(*x,j*): = *g*_2_(*x,j*:):= 0 for *j* = m.

Proof. Both *g*_1_ and *g*_2_ are non-increasing functions in *x* and *y*, hence properties 1 and 2 of De nition 3 are satisfied. Property 3 is satisfied because for all *y* 2 [m 1]

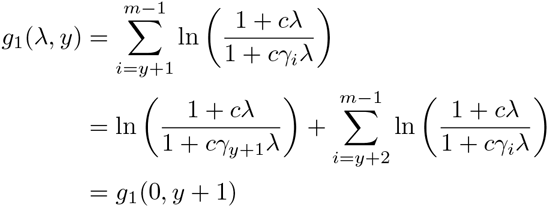

and

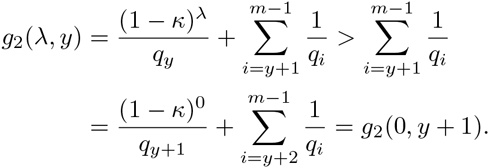

Drift analysis [30,32] is an important tool in runtime analysis of randomised search heuristics. Here we introduce a variant of the additive drift theorem [32] with its proof.

In the following, “(*x*_t+1_*x*_t_ +); t < T_a_” is the short-hand notation for “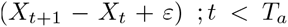” (see page 49 in [35]). Whenever we write an equality or inequality involving conditional expectation w.r.t. a σ-algebra, (e.g. 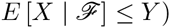, we have the “almost surely” meaning in mind.

**Lemma 22** (Additive drift theorem). Let 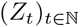 be a discrete-time stochastic process in [0; 1) adapted to any filtration 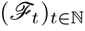. For any a ≥ 0, define 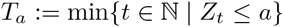. If for some ε > 0

1. 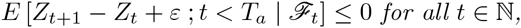

2. 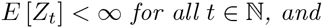

3. 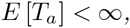
then 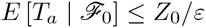.

Proof. Define the stopped process 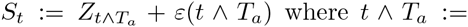 min(*t; T_a_*). By the definition of this process, it holds for all *t ε ℕ* almost surely that

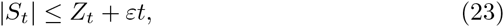

and, hence by condition 2 and 3, that for all 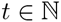,

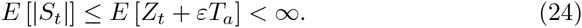

Also, by the definition of the process, for all t ∈ ℝ it holds in the case t *T* ≥ _a_ that,

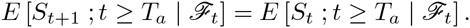

Furthermore, for all 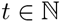, it holds in the case t < T_a_,

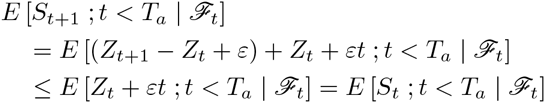

where the inequality is due to condition 1. Combining both cases, we have for all 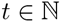,

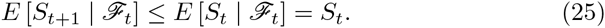

By (24) and (25), *S_t_* is a super-martingale, implying that for all 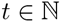,

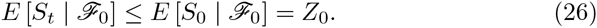

By (23) and (24), the dominated convergence theorem (see e.g. [35]) applies, and we get by (26)

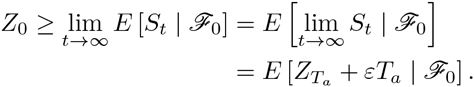

By noting that Z_Ta_ ≥ 0, the proof is now complete.

**Lemma 23** (Improved version of Lemma 5 in [16]). If *x* Bin ~ (λ, p) with p ≥ (*i*/λ)(1 + δ) and *i* ≥ 1 for some δ ∈ (0,1], then

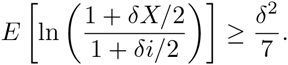

This improvement is due to the following generalisation of the lower bound in Lemma 17.

**Lemma 24.** For any *z* > 0, and all *x* ≥ 0 we have that

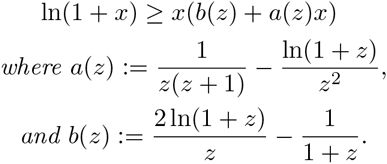

Proof. For *x* = 0, the result trivially holds. It then suffices to show that for all *x* ∈ (0, ∞)

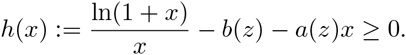

Note that *h(z)* = 0 and *h^'^(x)* = a(*x*) - *a(z)*. It follows from ln(1 + *x*) > 2x/(*x* + 2) for *x* > 0 (see (3) in [55]) that

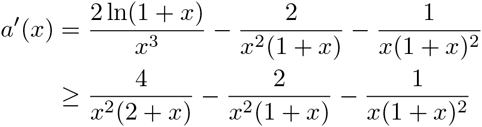

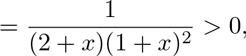

thus *a(x)* is an increasing function.

We separate two cases: for *x* ∈ (0,z], we have *a(x)* ≤ *a(z)* and *h^'^(x)* d 0, thus *h(x)* is decreasing on *(0,z] and h(x)* ≥ *h(z)* = 0; for *x* ∈ [*z*, ∞) we have h^'^(x) = *a(x)* - *a(z)* ≥ 0, *h(x)* is increasing on [z,∞) and *h(x) ≥ h(z)* = 0. We have shown that *h(x)* ≥ 0 for *x* > 0.

Note that the bound is tight at both *x* = 0 and *x* = z. The lemma does not cover the case *z* = 0, however at the limit, we get lim_z→0+_ b(z) = 1 and lim_z→0+_ *a(z)* = −1/2, and that corresponds to the bound given by Lemma 17.

**Corollary 25.** Let *x* ~ Bin(n,p) and μ:= E [X], then it holds that for all c > 0

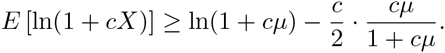

Proof. For p = 0 (or μ = = 0), the bound is trivial. Otherwise, for p > 0, applying Lemma 24 with *z* = cμ gives ln(1 + cX) ≥ b(cμ)cX + a(cμ)(cX)^2^, hence

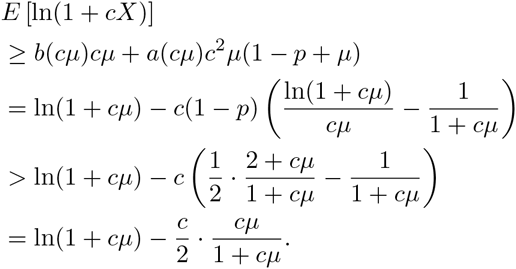

The last inequality is due to 1 - p < 1 and ln(1 + x)/x < (1/2)(x + 2)/(x + 1) for *x* > 0 (see (3) in [55]).

We now give the formal proof of Lemma 23.

Proof of Lemma 23. Let *y* ~ Bin(λ, (1 + δ)i/λ), then *y* ≼ X. Therefore,

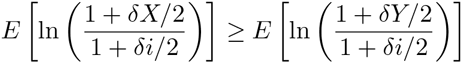

and it is sufficient to show that 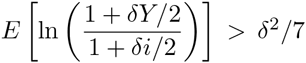 to complete the proof.

It follows from Corollary 25 (choosing c = =2) that

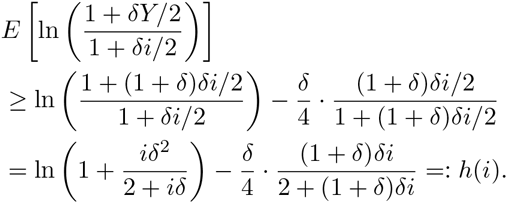

For all δ>0 and i≥1, it holds that

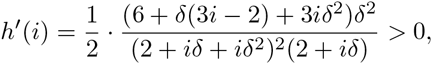

or h(i) monotonically increases in i.

Define r(δ): = 12 + 8δ + 3 δ^2^ + δ^3^ 2 ^4^ and s(δ): = (2 +δ)^2^(2 + δ + δ^2^) > 0, we get

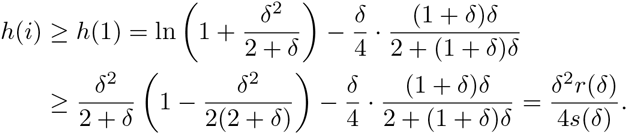

The last inequality is due to Lemma 17. We notice that 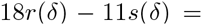 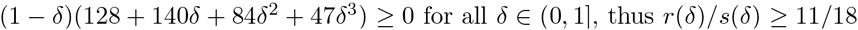 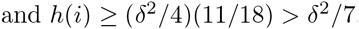.

**Lemma 26** (Lemma 6 in [16]). If *x* ~ Bin(λ, p) with p ≥(i/λ)(1 + λ), then E [e ^-kX^] ≤ e ^-ki^ for any *k* ∈ (0,λ).

Proof. The value of the moment generating function M_X_(t) of the binomially distributed variable *x* at t =-k is

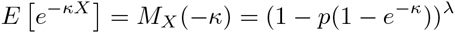

It follows from By Lemma 18 and from 1 + *κ* < 1 + that

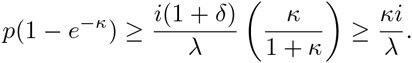

Altogether, we get E[e^-kX^]≤(1 - κi/λ)^λ^e^-ki^.

**Lemma 27.** Let {*x*_i_}_i∈[λ]_ be i.i.d. random variables, define 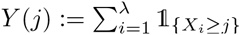 for any *j* ∈ ℝ. It holds for any a, b, c, *j* ∈ ℝ with c ≥ 0 and b ≤ λ that

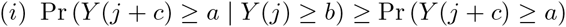

and for any non-decreasing function f

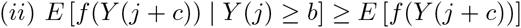

provided that both expectations are well-defined.

Proof. Define p: = Pr (*x*_i_ ≥ j) and q: = Pr (*x*_i_ ≥ *j* + c). For b ≤ 0 or p = 0, the result trivially holds. For b ∈ 2 (0,λ] and p ∈ (0,1], we have that q^'^: = Pr (*x*_i_ ≥ *j* + c | *x*_i_ ≥ j) = q/p ≥ q. Event *y* (j) ≥ b implies the existence of a set A ⊆ [λ] such that |A| ≥ |b| and *x*_i_ ≥ *j* for all i ∈ A. Define 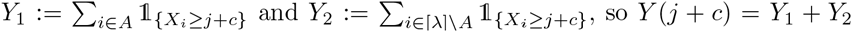. Clearly, conditioned on 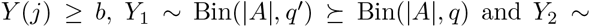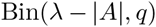. Therefore, the distribution of *y* (*j + c*) conditioned on *y* (j) ≥ *b* stochastically dominates Bin(|*A*|, *q*) + Bin(|*A*|,*q*) = Bin(λ,*q*), which is the (unconditional or original) distribution of *y* (*j* + *c*), and part (i) follows.

For part (ii), let *F*_1_(*x*): = Pr (*f(Y (j + c*)) < *x* | *y* (*j*) ≥ *b*) and *F*_2_(*x*): = Pr (*f*(*Y* (*j + c*)) < *x*); i.e. F_1_ and F_2_ are the conditional and the unconditional distribution functions of *f*(*Y* (*j+c*)) respectively. Then from part (i) we conclude that *F*_1_(*x*) ≤ F_2_(*x*) for any 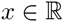, and by the properties of expectation,

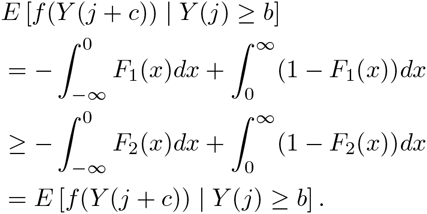

## Appendix C

The following results are used to analyse the tightness of the level-based theorem, i. e. Theorem 16.

Lemma 28 (Additive drift theorem (lower bound)). Let 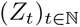 be a discrete-time stochastic process in [0,∞) adapted to any filtration 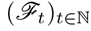. For any a ≥ 0, define 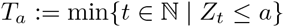. If for some ε > 0

1. 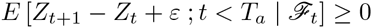 *for all* 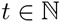, and

2. E [Z_t_] < ∞ for all 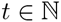.

3. E [T_a_] < ∞, then 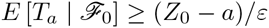.

Proof. The proof is similar to that of Lemma 22, i. e. starting by de ning the same stopped process *S_t_*. However, because the directions of the inequalities are inverted so *S_t_* is a sub-martingale, and in the end we overestimate *x*_Ta_ by a.

**Lemma 29.** If *x* ~ Bin(n,p) where *p* > 0, then for all 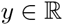

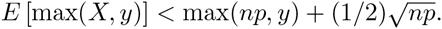

Proof. By Jensen’s inequality w. r. t. the square root, we have

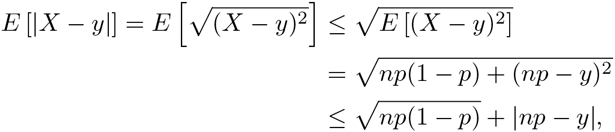

where the last inequality uses 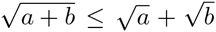 for a, b ≥ 0. Therefore, it holds that

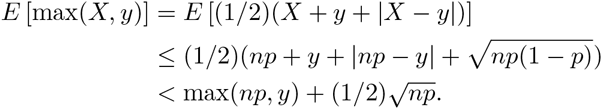

**Lemma 30.** If *x* ~ Bin(n,p) where p > 0 then it holds that E [X | *x* > 0] np + 1.

Proof. By definition,

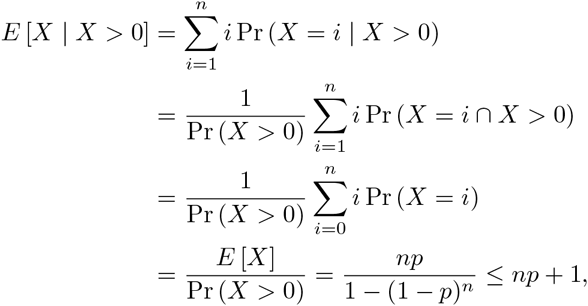

where the last inequality follows from Lemma 18.

## Footnotes

1 The first level can be *A*_0_ instead of *A*_1_ for some functions but that does not matter as far as we compute the sums correctly later on.

